# An active neural mechanism for relational learning and fast knowledge reassembly

**DOI:** 10.1101/2023.07.27.550739

**Authors:** Thomas Miconi, Kenneth Kay

## Abstract

How do we gain general insights from limited novel experiences? Humans and animals have a striking ability to learn relationships between experienced items, enabling efficient generalization and rapid assimilation of new information. One fundamental instance of such relational learning is transitive inference (learn *A>B* and *B>C*, infer *A>C*), which can be quickly and globally reorganized upon learning a new item (learn *A>B>C* and *D>E>F*, then *C>D*, and infer *B>E*). Despite considerable study, neural mechanisms of transitive inference and fast reassembly of existing knowledge remain elusive. Here we adopt a meta-learning (“learning-to-learn”) approach. We train artificial neural networks, endowed with synaptic plasticity and neuromodulation, to be able to learn novel orderings of arbitrary stimuli from repeated presentation of stimulus pairs. We then obtain a complete mechanistic understanding of this discovered neural learning algorithm. Remarkably, this learning involves active cognition: items from previous trials are selectively reinstated in working memory, enabling delayed, self-generated learning and knowledge reassembly. These findings identify a new mechanism for relational learning and insight, suggest new interpretations of neural activity in cognitive tasks, and highlight a novel approach to discovering neural mechanisms capable of supporting cognitive behaviors.

## 1 Introduction

Humans and animals can build representations of the world that allow them to generalize and rapidly gain insight from remarkably limited experience, yet how these abilities are implemented in the brain remains unclear [Tenenbaum et al., 2011, Tervo et al., 2016, Lake et al., 2017]. A fundamental basis of these abilities is the learning of *relations* between different experiences, embedding them in coherent structures that support generalizing inferences [Halford et al., 2010]. Through such “relational learning”, subjects are able to construct and also rapidly modify internal models of the world — variously referred to as “relational memory”, “schemas”, “cognitive maps” — that are increasingly recognized as essential to cognition [Eichenbaum, 2017, Behrens et al., 2018].

An classic instance of relational learning is transitive inference, that is, the ability to infer an ordered relationship between items not previously observed together. For example, after learning pairwise ordering between pairs of stimuli immediately adjacent in an ordered series (*A>B, B>C, C>D*, etc.), subjects immediately exhibit high performance on non-adjacent pairs not seen during previous training (*A>C, B>D*, etc.) Transitive inference has been observed in many species including humans, monkeys, rodents and pigeons (see Vasconcelos [2008], Jensen et al. [2019] for reviews).

Remarkably, animals and humans can also quickly combine previously learned orderings into a single overall ordering, after learning the ordering of their linking pair (“list-linking”: learning *A > B > C*, then *D > E > F*, then *C > D*, and inferring *B > E*) [Treichler and Van Tilburg, 1996, Nelli et al., 2023]. This striking ability for fast “reassembly” of existing knowledge [Nelli et al., 2023] implies that animals can quickly re-arrange representations of previously learned stimuli in response to limited novel information that does not explicitly include these previously learned stimuli.

Numerous models have been proposed to explain transitive inference. However, currently no known neural model explains the various results reported in behavioral experiments (see [Jensen et al., 2019] for a review). As a result, the neural mechanisms of transitive inference and knowledge assembly in relational learning remain elusive.

To investigate potential brain mechanisms of transitive inference, it is also possible to train artificial neural networks on a given ordered list, and observe their behavior [De Lillo et al., 2001, Kay et al., 2023, Nelli et al., 2023]. Trained networks often exhibit transitive inference over the learned lists, and their neural dynamics and representational geometry can be analyzed directly. However, because these lists are learned through an external, hand-designed optimization algorithm, these modelling approaches are intrinsically limited in clarifying how this learning itself occurs in the brain.

To address this challenge, here we take a different approach: rather than designing a learning algorithm ourselves, or training a neural network to correctly learn one particular series, we instead *meta-train* a neural network that can learn arbitrary new ordered lists. Building upon previous work in meta-learning, or “learning-to-learn” [Miconi, 2016, Miconi et al., 2018, 2019], we train neural networks endowed with biologically plausible (Hebbian) synaptic plasticity and neuromodulation. These networks can autonomously modify their own connectivity in response to ongoing stimuli and rewards. We train such networks specifically to be able to learn new ordered lists of arbitrary items from pairwise presentations, in settings similar to standard transitive inference experiments. Thus, the learning algorithm that these trained networks employ is unspecified *a priori* and entirely ‘discovered’ by the meta-training process.

We found that the resulting networks can learn new orderings from exposure to stimuli pairs and rewards alone, and exhibit transitive inference. Remarkably, despite not being explicitly trained to do so, they can also perform list-linking: after learning their linking pair, they can quickly reassemble separately learned lists.

In subsequent analyses, we obtained a complete mechanistic understanding of how these networks function, explaining how stimuli are represented by the network, how these representations are learned, and how transitive inference naturally emerges from these learned representations. Surprisingly, we show that fast reconfiguration of existing knowledge occurs through an active recall operation: during each trial, the networks actively and selectively *reinstate* representations of stimuli previously paired with those actually observed in the current trial. This allows learning (via synaptic plasticity) to alter the representation (and thus the list ranking) of these reinstated stimuli, in addition to the stimuli shown in the current trial. This active, cognitive process constitutes a novel neural mechanism for quickly reassembling existing knowledge in the face of new information, links synaptic mechanisms to cognitive and behavioral processes, and offers suggestions for investigating and interpreting neural data from experiments.

## 2. Methods

### 2.1 Terminology

Meta-learning, or “learning-to-learn”, consists in training an agent so that it becomes able to solve new instances of a general learning problem [Schmidhuber, 1987, Thrun and Pratt, 1998, Wang et al., 2016, Wang et al., 2018]. That is, instead of training the agent separately on each new instance of the problem, we *meta-train* a self-contained learning agent, so that it acquires the ability to learn autonomously and efficiently any given instance of the problem, including new instances never seen during training. Typical meta-learning problems in the literature include maze-solving [Duan et al., 2016, Miconi et al., 2018], bandit tasks [Wang et al., 2016], fast association between stimuli and labels [Santoro et al., 2016, Wang et al., 2016, Miconi et al., 2018], item value discovery problems such as the Harlow meta-task [Harlow, 1949, Wang et al., 2018], etc. Here we apply a meta-learning framework to plastic neural networks, training them to be able to learn new ordered series of arbitrary stimuli from pairwise presentations, as in classic transitive inference experiments.

In the following, to avoid confusion, we will maintain separate usage for the words *learning* and *training*. We use the word *learning* to denote within-episode learning of one particular ordered list, driven by synaptic plasticity in plastic weights. In contrast, we use the word *training* to denote the gradient-based optimization that occurs between episodes, and affects the structural parameters of the network, with the goal to improve within-episode plasticity-driven learning. This distinction follows that between “inner loop” and “outer loop”, respectively, in standard meta-learning terminology [Wang et al., 2016].

In each trial, after the network produces a response, it is given a binary feedback signal *R*(*t*) that indicates whether this response was correct or not. We call this signal “reward”, although we stress that it is merely an additional input to the network, and has no pre-defined effect on network structure by itself.

Finally, we use “stimulus” and “item” interchangeably.

### 2.2 Overview

We meta-train a recurrent neural network, endowed with synaptic plasticity and neuromodulation, to be able to autonomously learn an arbitrary serial order for a set of abstract stimuli, over the course of several pairwise trials. The general structure of the network and learning process largely follows Miconi et al. [2019].

All code for the following experiments is available online^1^.

The experiment is organized in successive *episodes*.

In each episode, the agent is tasked with learning a completely new ordering of randomly generated abstract stimuli (Fig. 1). The stimuli are randomly generated binary vectors of size 15; the number of items in the list to be learned varies from episode to episode between 4 and 9 (inclusive). Each episode consists of 30 *trials*, where each trial consists of the presentation of two stimuli (simultaneously), a binary response by the agent (“Choose Stimulus 1 or Stimulus 2”), and a binary feedback signal *R*(*t*) indicating whether the response was correct or not (that is, whether the chosen stimulus was in fact the higher-ranking of the two in the overall ordering).

**Figure 1.**
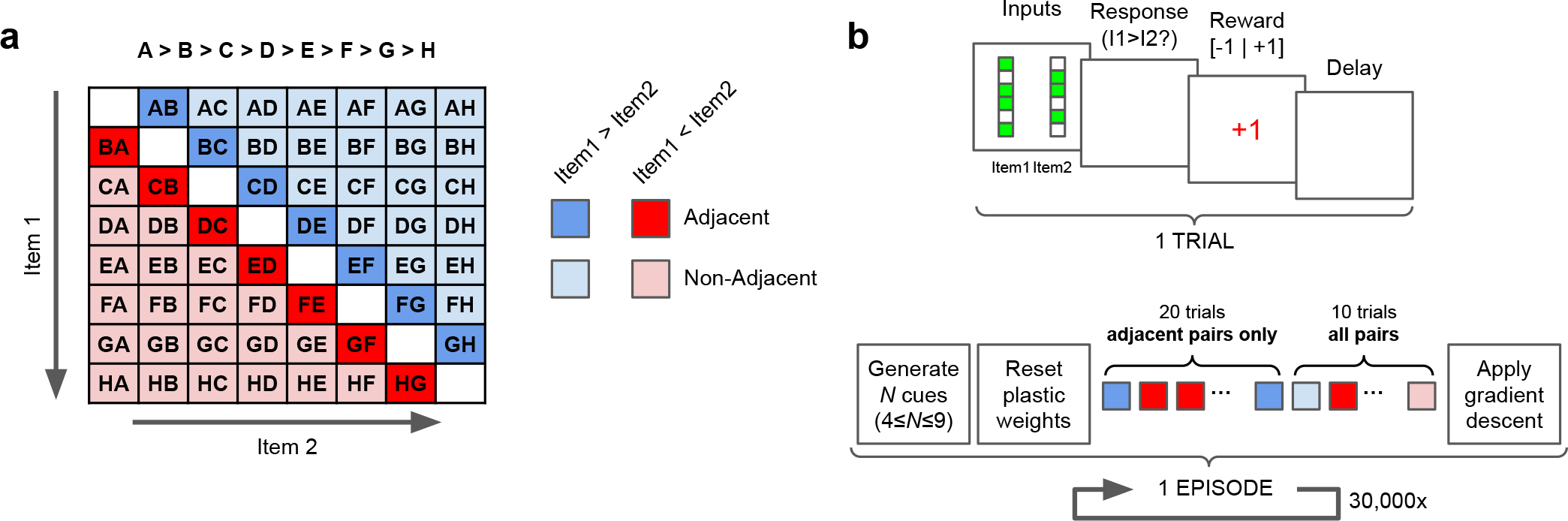
Schematic description of the transitive inference task. **a**: Possible stimulus pairs, colored according to adjacency and within-pair ranking. **b**: Each trial comprises a stimulus presentation, a network response, a feedback(reward) signal, and a delay. Each episode consists of 20 trials with only adjacent pairs, followed by 10 trials with all possible pairs. Plastic weights are reset between episodes.

The first 20 trials of an episode include only *adjacent* pairs, that is, pairs of stimuli with adjacent ranks in the series. The last 10 trials include all possible pairs (excluding identical pairs such as *AA* or *BB*), unless specified otherwise. Performance in a given episode is assessed as the proportion of correct responses over the last 10 trials.

**Figure 2.**
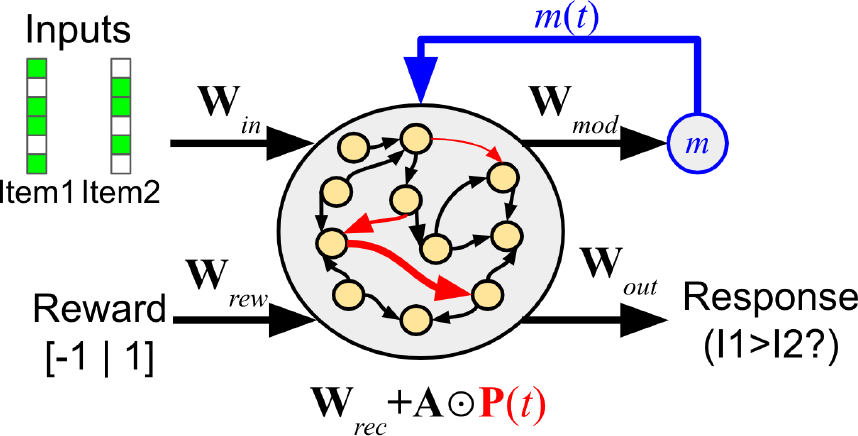
Organization of the network. Recurrent connections are plastic and change over the course of an episode.

Within each episode, the agent undergoes synaptic changes as a result of ongoing synaptic plasticity, gated by a self-generated modulatory signal (see below). From these synaptic changes, a successful agent will learn the correct order of all stimuli over the course of the episode. To generate such agents, after each episode, we apply gradient descent to the structural parameters of the network (the base weights and plasticity parameters, as well several other parameters; see below), in order to improve within-episode, plasticity-driven learning. The loss optimized by gradient descent is the total reward obtained over the whole episode, with rewards in the last 10 episodes being up-weighted by a factor of 4 (this up-weighting only affects the meta-learning loss for gradient descent and does not affect the reward signal actually perceived by the network).

The gradients are computed by a simple reinforcement learning (RL) algorithm, namely Advantage Actor Critic (A2C) [Mnih et al., 2016], which is well understood and can be interpreted as a model of dopamine-based learning in the brain [Wang et al., 2018]. See section B for details.

### 2.3 Network organization

The network is a recurrent neural network containing *N* = 200 neurons. The inputs **i**(*t*) consist of a vector that concatenates the stimuli for the current time step, the reward signal *R*(*t*) for the current time step (0, 1 or −1), and the response given at the previous time step (if any), in accordance with common meta-learning practice [Wang et al., 2016, Duan et al., 2016]. The network’s output is a probability distribution over the two possible responses (“Choose Stimulus 1” or “Choose Stimulus 2’). At response time, the agent’s actual response for this trial is sampled from this output distribution; at other times the outputs are ignored.

Recurrent connections within the network undergo synaptic changes over the course of an episode, modelled as simple neuromodulated Hebbian plasticity as described in Section 2.4.

Formally, the fully-connected recurrent network acts according to the following equations:

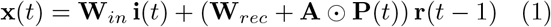

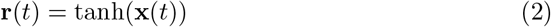

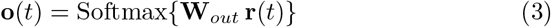

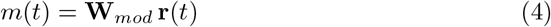

Here **i**(*t*) is the vector of inputs, **x**(*t*) is the vector of neural activations (the linear product of inputs by weights), **r**(*t*) is the neural firing rates (activations passed through a nonlinearity), **o**(*t*) is the output of the network (i.e. the probability distribution over the two possible responses), and *m*(*t*) is the neuromodulatory output that gates in plastic changes (see below).

**W**_*rec*_ and **A** (capital *α*) are the base weights and plasticity parameters (plasticity learning rates) of the recurrent connections, respectively. They are *structural* parameters of the network and do not change during an episode, but rather are slowly optimized between episodes by gradient descent. By contrast, **P**(*t*) is the *plastic* component of the weights, reset to 0 at the start of each episode, and changing over an episode according to the plasticity rule described below. *⊙* represents the pointwise (Hadamard) product of two matrices.

Note that these equations are simply the standard recurrent neural network equations, except that the total recurrent weights are the sum of base weights **W**_*rec*_ and plastic weights **P**(*t*) multiplied by the plasticity parameters **A**.

Crucially, within an episode, only the plastic recurrent weights **P**(*t*) are updated according to the plasticity rule described below. All other parameters (**W**_*in*_, **W**_*rec*_, **W**_*out*_, **W**_*mod*_, **A**) are fixed and unchanging within an episode, but are optimized by gradient descent between episodes.

### 2.4 Synaptic plasticity

Each connection maintains a so-called Hebbian eligibility trace **H**(*t*), which is a running decaying average of the product of outputs and inputs:

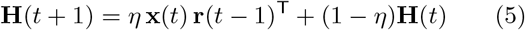

Note that, because the network is recurrent, **x**(*t*) constitutes the “outputs” while **r**(*t−* 1) constitutes the “inputs” at any given connection (cf. Eq. 1).

Finally, the network continually produces a neuromodulation signal *m*(*t*) that gates the Hebbian trace into the actual plastic weights **P**:

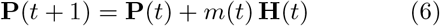

We emphasize that **P**(*t*) is initialized to 0 at the beginning of episodes and changes according to the above equations (without any reinitialization) over the course of an entire episode. Note that while **H**(*t*) is decaying, **P**(*t*) does not decay within an episode.

In the equations above, *η* is a parameter of the network, optimized by gradient descent between episodes. Importantly, across many experiments, we observed that training consistently settles on a value of *η≈* 1. This means that H(t) is almost fully updated at each time step, with essentially no memory of past events prior to time *t−* 2. As such, we can approximate the actual plasticity process as follows:

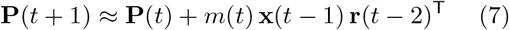

This observation plays an important role in the elucidation of network behavior, as described below.

### 2.5 Experimental settings

Our objective is not simply to train a successful learning network, but also to fully understand the neural mechanism of learning in this trained network. To this end, we deliberately sought to simplify trial structure as much as possible, while retaining the essential structure of real-world experiments.

First, we restricted each trial to the shortest possible duration that still allowed for successful training, which we found to be exactly four time steps: stimulus presentation on step 1, network response at step 2, external feedback (reward) at step 3, and lastly a delay time step before the start of the next trial, as step 4.

Second, we reset neural activations **x**(**t**) and **r**(**t**) and Hebbian eligibility traces **H**(*t*) at the start of each trial (but not plastic weights **P**(*t*)). This is to ensure that neural dynamics remain confined to a single trial, and that only plastic weights can carry memory of past events from one trial to the next^2^.

## 3. Results

### 3.1 Two different algorithms for relational learning

Across multiple runs, the meta-training procedure described above consistently generates a high-performing learning agent (Fig. 3). Inspection shows that all runs that reach the higher performance level seem to follow a similar strategy, which is analyzed below. Interestingly, some runs reach a slightly lower performance point, which turns out to reflect a different, less efficient strategy. Importantly, these two generic strategies also differ in that only the high-performing one is able to perform list-linking, as explained below. These two discrete outcomes appear consistently across various experimental settings, though the proportion of each varies (see section 5).

**Figure 3.**
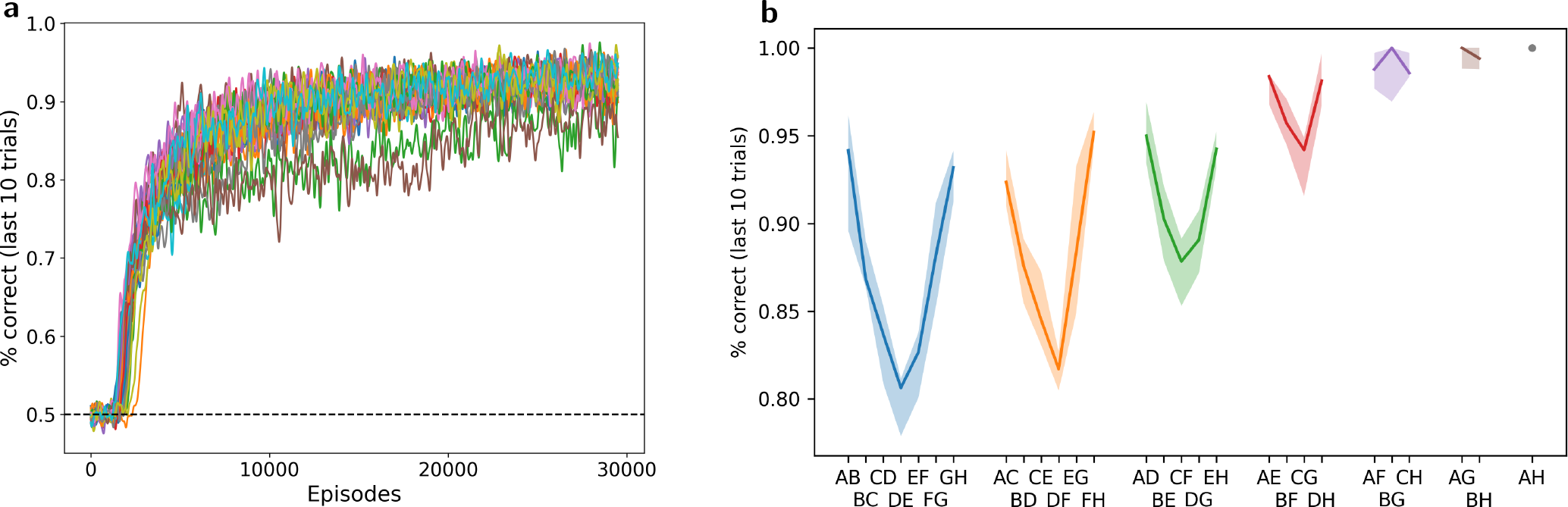
Transitive inference and realistic behavioral patterns. **a**: Meta-training loss: Mean performance in the last 10 trials of each episode, for 30000 episodes of meta-training (30 separate runs, all curves smoothed with a boxcar filter of width 10). Notice the two slightly lower curves, indicating two runs settling on the suboptimal solution (see Section 5). **b**: Test performance: mean performance in the last 10 trials of 2000 test episodes (with the same frozen network but different randomly generated stimuli), for each pair, arranged by “symbolic” (intra-pair) distance. More distant pairs elicit higher performance (symbolic distance effect), as observed in animal and human experiments.

All following results were obtained from a single network representative of the high-performance strategy, unless stated otherwise.

### 3.2 Transitive inference, symbolic distance effect and end-anchor effect

To assess the behavior of the trained network, we ran 2000 copies of the fully trained network on one full episode of 30 trials, each with a different randomly generated set of 8 stimuli (from *A* to *H* inclusive) and a randomly chosen pair for each trial. Fig. 3B reports performance in the last 10 trials of this test episode, separately for each pair, with pairs arranged according to symbolic distance (i.e. the absolute difference in rank between both items in the pair).

These plots show two strong effects typical of transitive inference in animal and human experiments [Vasconcelos, 2008, Jensen et al., 2019]. First, we observe the so-called *symbolic distance effect* : performance is higher for the pairs with the highest intra-pair distance in rank. In particular, performance is lowest for adjacent pairs (AB, BC, etc.). This is remarkable, since the first 20 trials involved adjacent pairs only (as in animal experiments): performance is worse on the “training set”.

Second, we observe an *end-anchor effect* (also known as “serial position effect”): performance is consistently higher for item pairs that involve the highest and lowest ranked items, rather than “interior” pairs of comparable symbolic distance (e.g. performance on *AC* and *FH* is higher than performance on *CE* or *DF*) This effect is seen on the U-shape of the curves in Fig. 3B, similar to that observed in animal and human data [Vasconcelos, 2008, Jensen et al., 2019]. Thus, the trained network successfully demonstrates transitive inference, and reproduces behavioral patterns consistently observed in animal and human experiments.

Notably, in additional experiments, we found that the network is robust to so-called “massed presentation” of one single pair, which is known to disrupt certain learning models of transitive inference [Jensen et al., 2019, Lazareva and Wasserman, 2012] (see Section G for details).

### 3.3 List-linking: fast reassembly of existing knowledge

Monkeys and humans can quickly link together separate orderings after learning the correct connecting pair. That is, after learning *A > B > C > D* and *E > F > G > H* separately, and then learning *D > E*, they can quickly infer ordering across the whole joint sequence (*C > F, B > G*, etc.) [Treichler and Van Tilburg, 1996, Nelli et al., 2023]. This *list-linking* ability implies that the presentation of a pair can affect the subjective ranking not just of the shown pair itself, but also of other elements not shown in the current trial.

We ran the trained networks on 10 trials using adjacent pairs from *ABCD*, then 10 trials using adjacent pairs from *EFGH*, and then finally 4 trials with *D* and *E*. Then we estimated performance on one single trial, which could use any pair from the whole *ABCDEFGH* sequence. This was done over 2000 runs, again with different randomly generated stimuli for each run. Results for the last, “test” trial are shown in Fig. 4. Examinations of results for pairs including items from both sub-lists (e.g. *CE, CF, BG*, etc.) confirms that the network successfully linked the two lists into a coherent sequence. We observe that performance is consistently poor for the pairs immediately adjacent to the linking pairs, especially from the earliest-learned sequence (here, *CD*); this effect is also reported in monkey experiments [Treichler and Van Tilburg, 1996, Table 2].

**Figure 4.**
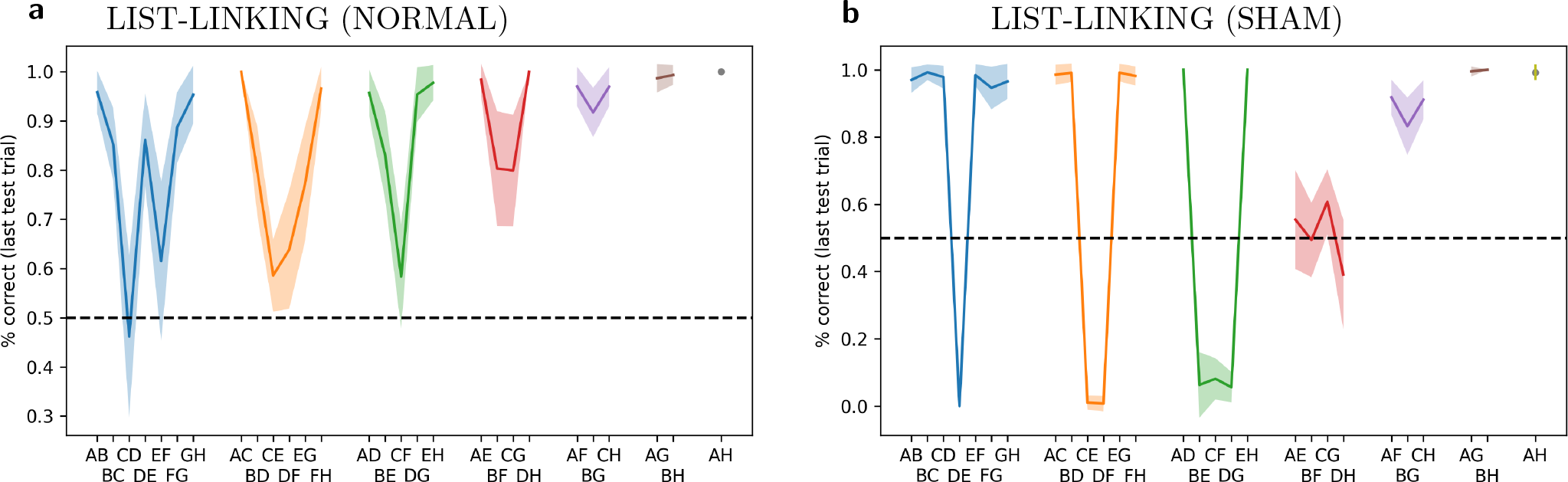
Networks successfully perform list-linking. **a**: Performance for each pair in a list-linking setting, learning *ABCD*, then *EFGH*, then *DE*, then testing on a single trial with any pair. Performance shown for last, test trial only. **b**: “Sham” list-linking: last training phase on *EF* (non-linking pair) rather than *DE*, for one trial only.

To further assess this ability, we ran a control experiment with “sham” list linking. This was exactly identical to the one described above, except that instead of 4 trials on the linking pair *DE*, we showed only 1 trial with a non-linking pair *EF*. Thus, no linking could possibly occur: the network had no information about which of the two sub-lists was supposed to come first in a global ordering. We findperformance is considerably *below* chance on neighbouring pairs that run across lists, but very high on more distant pairs(Fig. 4B). These results suggest that the network represents rank in a way that transfers across lists, and can compare ranks between items of different lists (i.e. because *E* is “high” in the *E > F > G > H* sequence, while *D* is “low” in the *A > B > C > D* sequence, the system infers that *E > D*). This cross-list transfer is also observed in animal experiments [Jensen et al., 2019].

## 4 Neural mechanisms of learning

### 4.1 Overview of the discovered learning algorithm

The previous results show that the network can efficiently learn list order within and across lists from pairwise presentations. We now seek to understand how this learning takes place. Because the network is a learning network, this can be divided into two questions: first, how incoming information is represented and manipulated in network activity; and second, how these representations are learned by the network across trials.

As explained in detail over the following sections, the neural learning algorithm discovered by the meta-training process can be summarized as follows:

1. *Step 1: feedforward stimulus representations*. At step 1, the network perceives the two items for the current trial. Neural activations **r**(*t* = 1) reflect the feedforward representations of both items, as determined by the fixed, non-plastic input weights **W**_*in*_. Input weights for the two items turn out to be strongly negatively correlated with each other, and thus the resulting representation of the pair is essentially a *subtraction* between the feedfoward representations of both items in isolation.
2. *Step 2: Trial decision and item reinstatement (multiplexed)*. At step 2, recurrent (plastic) weights project the previous feedforward representations into *learned* representations (step-2 representations are learned because they are determined by plastic recurrent weights, which are modified by within-episode learning). These learned representations turn out to encode rank geometrically: stimulus rank is encoded by the alignment of the stimulus’ step-2 representation with a fixed “decision axis”, which turns out to be strongly correlated with the output weight vector **W**_*out*_. Because step-1 feedforward representations of pairs are subtractions between the representations of each item, the step-2 representation of the pair is also approximately the subtraction between the learned, step-2 representations of both items. The alignment of this subtraction with the decision axis (output weight vector) thus automatically encodes the rank difference between the items, and determines the network’s response for this trial, thereby explaining transitive inference (once adequate representations are learned for each item, subtraction can operate on any pair of items) and the symbolic distance effect (larger differences overcome the imprecision in each individual item’s representation). *In parallel to this*, the network also reinstates in its activity the (recoded) step-1 representations of both shown items, as well as other items previously “*coupled*” with these (see below).
3. *Step 3: Reward delivery and decision axis reinstatement*. At step 3, a feedback signal informs the network about response correctness. At this step, a moderate burst in *m*(*t*), of constant sign and independent of response or reward, induces Hebbian learning between **r**(*t* = 1) and **r**(*t* = 2). This plasticity “stamps in” the coupling between step-1 feedforward representations and step-2 reinstatements of their recoded versions discussed above. This ensures their “coupling” and thus future joint reinstatements at step 2 of future trials. *In parallel*, the network reinstates in its activations the (recoded) representation of the decision axis, with the appropriate sign depending on the network’s response for the current trial.
4. *Step 4: Representation learning*. At step 4, a larger, reward-sensitive burst of *m*(*t*) induces Hebbian learning (with the appropriate sign depending on response correctness) between **r**(*t* = 2) and **r**(*t* = 3). This links the reinstated, recoded step-1 representations of the shown items *as well as* their coupled neighbours (which were all reinstated at step 2, as mentioned above) to the reinstated, recoded representation of the decision axis (which was reinstated at step 3, as mentioned above). The resulting modulated Hebbian learning modifies plastic weights in such a way that the step-2 representations of all reinstated items (both currently shown items and not-shown neighbouring items coupled with these) will be shifted in the appropriate direction along the decision axis, ensuring that this alignment encodes stimulus rank.

This discovered neural algorithm learns ordered lists efficiently, in a way that automatically transfers to non-adjacent pairs (transitive inference), is more robust for pairs with larger difference (symbolic distance effect), and uses information from a given trial to adjust not just the items shown at this trial, but also their known neighbours in rank order (enabling list-linking by only showing the linking pair).

The latter ability, in particular, is the result of a cognitive act: the selective reinstatement in memory of items previously shown, in order to support delayed Hebbian learning for these reinstated items in addition to items shown in the current trial. This active reinstatement supports fast reassembly of existing knowledge upon presentation of the linking item alone. In contrast, the alternative discovered neural mechanism (which is passive and lacks this active reinstatement) fails the list-linking task (see Section 5).

We now proceed to explain in detail how the network operates and learns.

### 4.2 Neuromodulation dynamics

First, we examine the neuromodulatory output produced by the network over the course of an episode, and how it is affected by received reward signals. Fig. 5 shows the neuromodulatory output *m*(*t*) (red) and the delivered reward (blue) over a single full episode, in several runs with the same trained network (each with different randomly generated stimuli). Each spike in the blue (reward) curve corresponds to the reward (positive or negative) delivered at step 3 of one trial.

**Figure 5.**
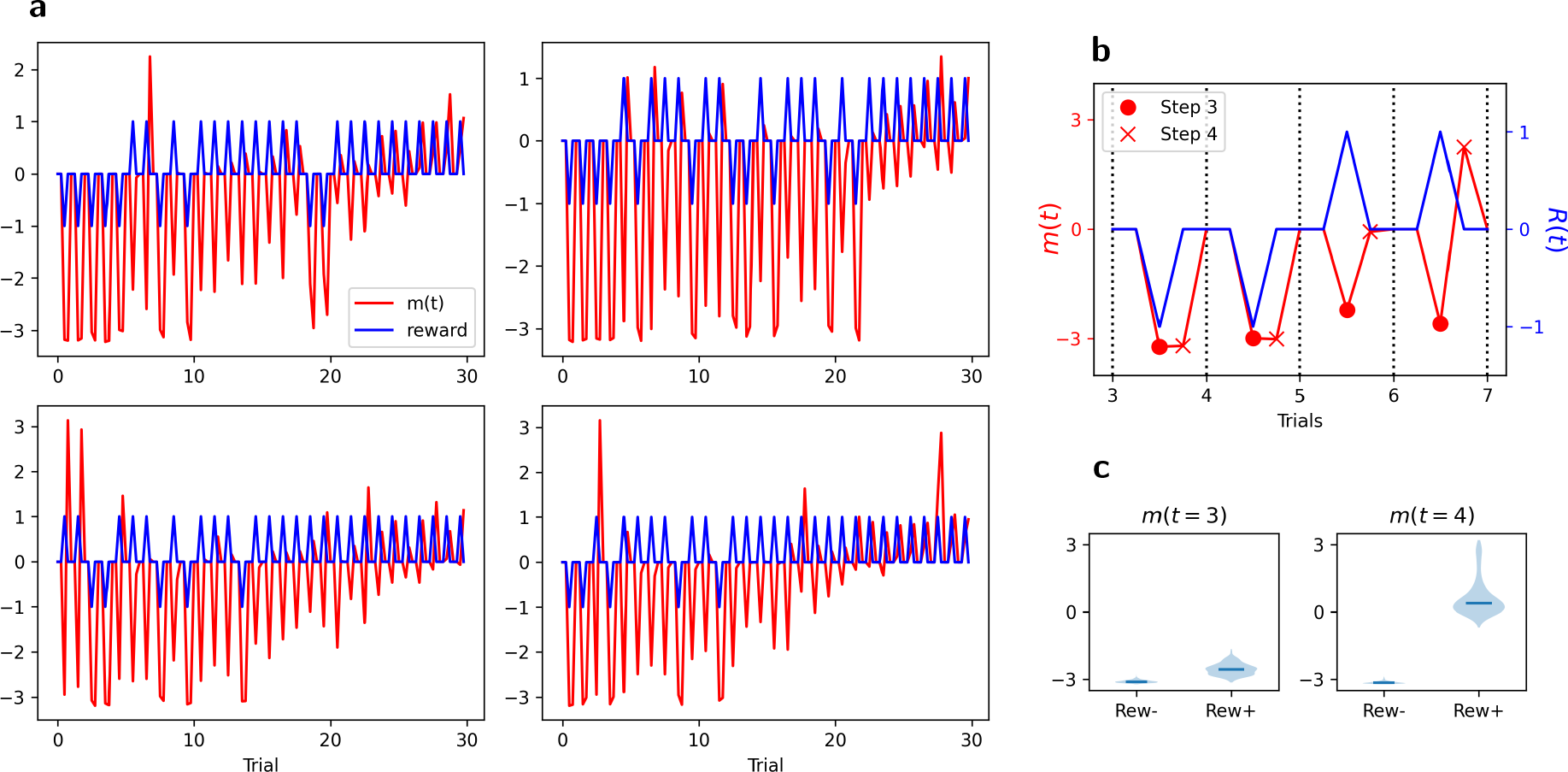
Dynamics of received reward signal and network-generated neuromodulatory output. **a**: Neuro-modulatory output *m*(*t*) (red) and reward (blue), for four different episodes (each with different stimuli but with the same initial trained network). *m*(*t*) is zeroed out for the first two time steps of each trial (see text). **b**: Zoom on trials 3-6 from first graph in **a**: In each trial, *m*(*t* = 3) (*•*) is consistently negative independently of reward sign, while *m*(*t* = 4) (*×*) is negative for negative rewards and positive or very small for positive reward. **c**: Violin plot of *m*(*t* = 3) and *m*(*t* = 4) for trial 5 across all 2000 runs, shown separately for correct and incorrect responses.

Note that the trace has been zeroed out for the first two steps of each trial, since no weight modification can occur at these times due to the reset of **x**(*t*) and **r**(*t*) at the start of each trial.

We first observe that *m*(*t* = 3) (at the time of reward delivery, coincident with blue spikes) appears not to be sensitive to rewards: it is consistently negative early in the episode, independently of reward received, and gradually increases across the episode. By contrast, *m*(*t* = 4) (time step just after the reward) is highly selective for reward. It is strongly negative for negative rewards and either positive or very small for positive rewards. This suggests that neuromodulatory outputs at time steps 3 and 4 of a given trial play different roles in learning the task.

### 4.3 Learned representations and decision axis

We again ran the trained network on a full episode of 30 trials, on 2000 runs with different randomly generated stimuli. We first focus on network activity at time step 2 of trial 20, that is, the “response” step of the last trial with adjacent-only pairs. We consider the vector of neural firing rates **r**(*t* = 2) at that point in time, for each of the 2000 runs, as a separate data point, and perform principal component analysis (PCA) over this data.

Fig. 6 plots all 2000 data points projected on the first two principal components of the data, colored according to various features of the corresponding trials. We first note that the first principal component (PC1) is dominant: it explains 27% of the variance, while the next PC explains 4%.

**Figure 6.**
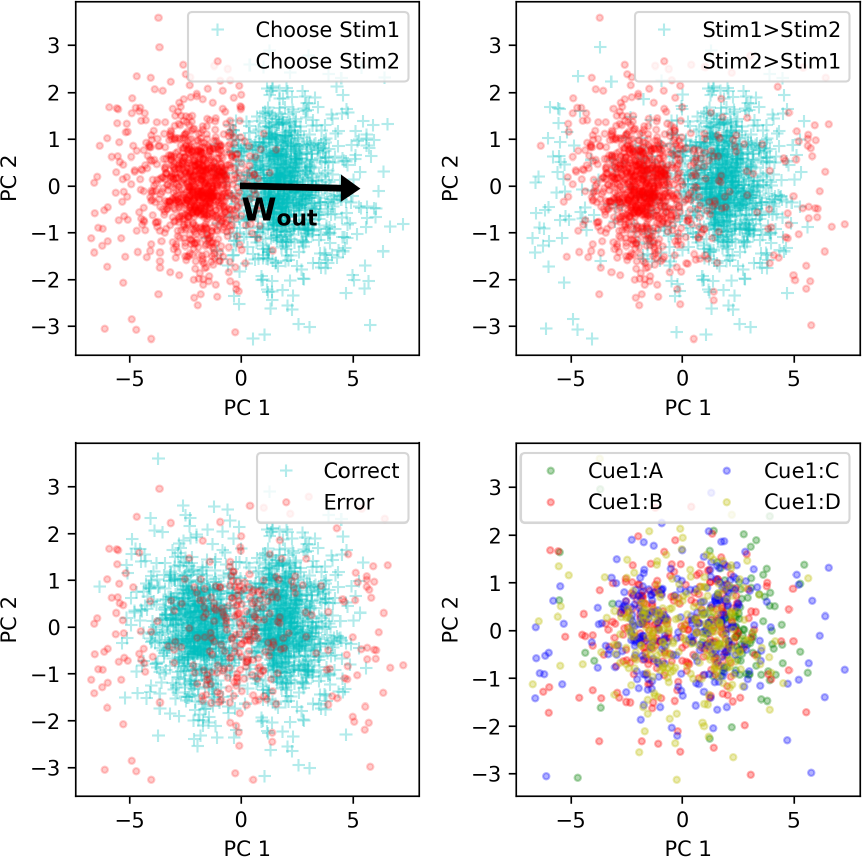
Principal component analysis of neural activity. Each data point represents **r**(*t*) at time step 2 of trial 20, for 2000 different runs. PC1 is associated with network response and is strongly aligned with the vector of output weights (top-left). Other graphs label data points according to whether the items were shown in the right order for this trial (top right), whether the response was correct (bottom left), and the identity of item 1 (bottom right; only 4 possible items shown for clarity)

We observe a strong gradient along PC1 between thew two possible responses (Fig. 6, top-left). This suggests that the direction of PC1 is relevant to the network’s response. What could cause this relevance? Recall that the output weight matrix **W**_*out*_ (trained by gradient descent between episodes, and fixed and unchanging during any episode) is a 2 *×N* matrix, projecting **r**(*t*) onto logits for each of the two possible decisions. We first observe that, unsurprisingly, the two rows of **W**_*out*_ are strongly anti-aligned, with correlation *r < −*.9. We therefore summarize **W**_*out*_ as a single vector **w**_*out*_, computed as the difference between the two rows (results are similar if we use either row instead).

We find that this output weight vector shows high alignment with PC1, with correlation *r >* .9. We plot the projection of this output vector over PC1 and PC2 in Fig. 6, top left, illustrating this high alignment with PC1. In other words, the learned, step-2 representations of input pairs (resulting from one step of recurrence through learned, plastic weights) are arranged along a “decision axis” aligned with the output weights of the network, immediately explaining the readout from these representations to the corresponding decision.

We note that at trial 20, this response is usually, but not always correct: other graphs on Fig. 6 show the same data colored according to whether the item pair was shown in the right order or not (top right), and according to whether or not the response was correct (bottom left). We note that most responses were indeed correct, except near the middle of the plot (presumably representing uncertainty from the network and stochasticity in the response) and, more surprisingly, at the extremes on either side, where a distinct “halo” of incorrect responses can be seen.

Inspection reveals that these extreme responses correspond to those runs in which trial 20 featured a middle pair (e.g. *DE* if the list to be learned had 8 items from *A* to *H*) which, fortuitously, was not seen before in the episode for this particular run. This led the network to assume that the higher-order member of the new pair (*DE*) was “low” (since it was at the end of *A > B > C > D*) while the lower-order member was “high” (*E > F > G > H*), leading to a confident wrong assessment when first seeing the new pair *DE*. This further confirms that PC1 represents a decision axis, with direction representing decision sign and distance representing confidence.

### 4.4 Representation of individual items and item pairs

How does the system align the learned representations of each item pair with the output weight vector, in a way that transfers to non-adjacent pairs? One possibility is that these learned representations at step 2 might contain information about the identity and rank of each individual item in the pair. However, this does not seem to be the case. Neural vectors don’t seem to be consistently separated by the rank of the first item (Fig. 6, bottom right). Furthermore, using various classification methods, we failed to reliably decode the rank of either the first or the second item from neural data at step 2 (Fig. S1).

To understand how pair representations are built, we first examine the representations of isolated single items. That is, we present each possible stimulus *X ∈{A, B, C, D, E, F, G,} H* to the network as item 1, in isolation (not paired with any other), and observe the resulting neural activity **r**(*t*) at time steps 1 and 2. When showing item *X* in isolation, we denote **r**(*t* = 1) with the symbol ***ψ***_*t*1_(*X*), and **r**(*t* = 2) with the symbol ***ψ***_*t*2_(*X*). Thus, ***ψ***_*t*1_(*X*) is the “feedforward”, step-1 representation of *X*, determined solely by the fixed input weights. By contrast, ***ψ***_*t*2_(*X*) is the “learned”, step-2 representation, produced by applying one step of recurrence through the learned, plastic weights to ***ψ***_*t*1_(*X*).

We freeze the plastic weights **P**(*t*) at trial 20, and apply these frozen networks to each individual item in isolation for two steps, which allows us to extract ***ψ***_*t*1_(*X*) and ***ψ***_*t*2_(*X*) for each item *X ∈ {A, B, C, D, E, F, G, H}*. We then compute the alignment (correlation) between ***ψ***_*t*2_(*X*) and the output weight vector **w**_*out*_, for each single item *X*. As shown in Fig. 7, we find that the step-2 neural representation of each single item ***ψ***_*t*2_(*X*) aligns with the output weight vector **w**_*out*_ proportionally to the item’s rank: ***ψ***_*t*2_(*A*) has strongly positive correlation with **w**_*out*_, ***ψ***_*t*2_(*H*) has strong negative correlation with **w**_*out*_, and intermediate items follow a monotonic progression (Fig. 7). In other words, the network somehow learns to map each individual item *X* (in isolation) to a learned representation ***ψ***_*t*2_(*X*), whose alignment with the decision axis encodes the item’s rank in the sequence.

**Figure 7.**
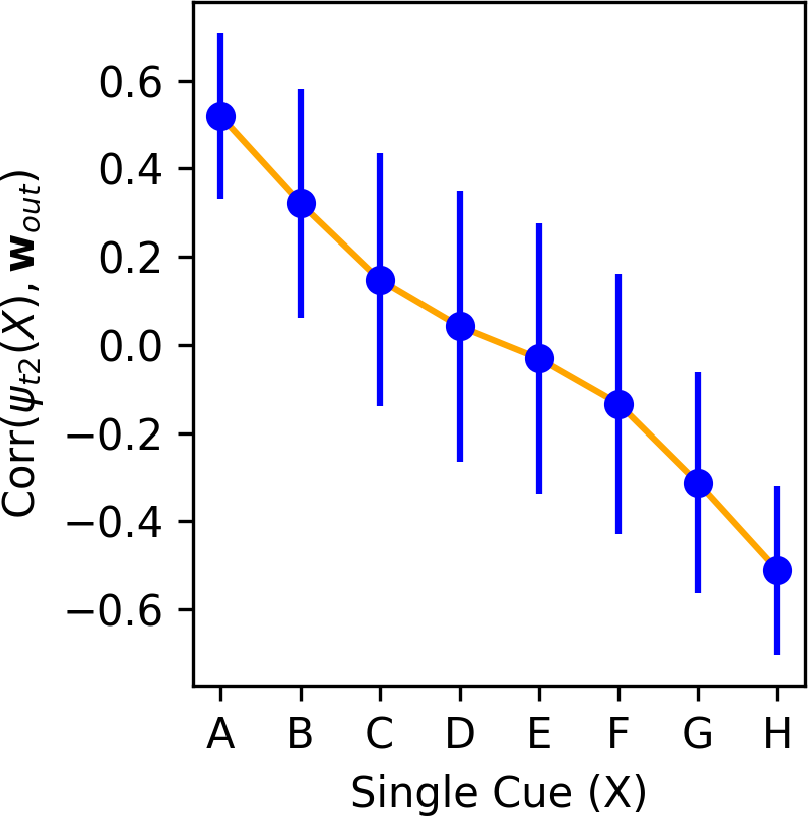
Correlation between the step-2 representation of each item ***ψ***_*t*2_(*X*) and the output weight vector **w**_*out*_, for each item *X ∈ { A, B, C, D, E, F, G, H,}* at trial 20. Mean and standard deviation over 2000 runs of one episode with a trained network, each with independently generated item stimuli.

In the actual task, stimuli are not presented in isolation, but in pairs. How are those learned representations of single items combined to represent pairs of items? Examining input weights **W**_*in*_, we find that the input weights for both items in the stimulus pair are strongly anti-correlated (*r ≈*.*−*9). As a result, a given item’s representation when it is shown as item 2 is essentially the negative of its representation when shown as item 1. Therefore, when a pair of items (*X, Y*) is presented as input, the network automatically computes a *subtraction* between the representations of both items (**r**(*t* = 1) *≈****ψ***_*t*1_(*X*) *−****ψ***_*t*1_(*Y*)). At the next time step, application recurrent weights transform this subtraction into ***ψ***_*t*2_(*X*) *−* ***ψ***_*t*2_(*Y*) (neglecting the nonlinearity). Since ***ψ***_*t*2_(*X*)’s alignment with the output weight vector **w**_*out*_ is proportional to its rank, the alignment of their subtraction with **w**_*out*_ is proportional to their difference in ranks. This allows the output weights to automatically produce an output proportional to the difference in rank between the items, which is precisely the expected behavior.

### 4.5 A simple representational scheme for transitive inference

These operations represent a simple, intuitive mechanism for transitive inference: once the correct representation ***ψ***_*t*2_(*X*) for each individual item *X* has been learned, the subtractive operation immediately generalizes to non-adjacent pairs. Furthermore, more distant pairs imply a larger difference in the projection points of either isolated item on the decision axis, overcoming the noise in the actual alignment of each individual item; this suffices to cause a symbolic distance effect, where more distant item pairs are more accurately judged.

Again, we can visualize a consequence of this subtractive operation in the “halo” of incorrect responses at the extremes in Fig. 6. In these runs, the system incorrectly assigns a very high and a very low order to each item, because its history of input presentations spuriously showed them as the last and first of two separate sub-sequences. As a result, the representations of these stimuli have extreme (and extremely wrong) projections along the decision axis, and their subtraction will also have a large projection along this axis (in the wrong direction), causing a “confidently wrong” response.

While this organization successfully explains order judgement and transitive inference, it assumes a correct learned representation for each single item, with rank-proportional alignments over the decision axis. This leaves open the question of how these representations are learned, which we address in the following sections.

### 4.6 Representation learning

#### 4.6.1 Simple Hebbian learning does not explain results

How does the network learn to map single items to rank-appropriate representations? Recall that each trial involves four steps: item presentation, then network response, then feedback, and lastly a delay. In addition, the network is endowed with Hebbian learning, where the weight change in a connection at time *t* is essentially the product of input and output at time *t−* 1, multiplied by a (signed) network-controlled *m*(*t*) signal.

In theory, this would suffice to adjust the step-2 representation of each item in the correct direction, based on feedback at time step 3, through simple reward-modulated Hebbian learning: that is, items presented at time step 1 would become associated with the response given at time step 2, and this association would be modulated positively or negatively based on feedback at time step 3, leading to a reinforcement of the correct association.

However, this cannot be the sole mechanism at play. Such a mechanism would only affect the representation of the items shown in the current trial. By contrast, network behavior (namely, the list-linking results in Fig. 4) shows that a item pair presentation can affect future responses to other items, not shown in the current trial. In other words, information about the rank order of the currently shown pair somehow reconfigures the rank of other items, not seen in the current trial. This is particularly surprising since we explicitly reset neural activity (but not plastic weights) at the start of each trial, preventing previous-trials information from echoing into the current trials through recurrent activity. How does this transfer of information happen?

#### 4.6.2 Step-by-step changes in plastic weights and learned representations

To address this question, we examine changes in the plastic weights (and the resulting representations) at *each successive time step*. We select runs in which the network sees the pair *DE* or *ED* at trial 20, but not in any previous trial (we focus on *DE* for illustration only; as shown in Fig. S2, similar results hold for other pairs). This presentation of a new pair of intermediate rank, not seen before, invariably causes an erroneous response in this trial. However, we want to observe the actual change resulting from this error.

For every step of trial 20, we freeze the current **P**(*t*) at this time step, and run the network with these frozen plastic weights for two time steps, presenting each stimulus *X ∈{A, B, C, D, E, F, G, H}* in isolation. This allows us to extract the current step-2 representation of each item ***ψ***_*t*2_(*X*), as encoded by the present plastic weights **P**(*t*) *at this time step*, and compute its correlation with the output weight vector **w**_*out*_ (that is, exactly the same quantity plotted in Fig. 7). By measuring this quantity successively at each time step, we can observe how each item’s learned step-2 representation changes its alignment with the decision axis, from one time step to the next.

Fig. 8 shows the change in alignment between the step-2 representation of each item and output weights, measured separately at each successive time step of trial 20 (when pair *DE* is shown for the first time). Note that, because we reset activations **x**(*t*) and **r**(*t*) at the start of each trial, no weight changes occur for the first two time steps of a trial (cf. Eq. 5). We observe that step 3 (when the reward signal is delivered to the network) produces only small changes, restricted to items *D* and *E*.

**Figure 8.**
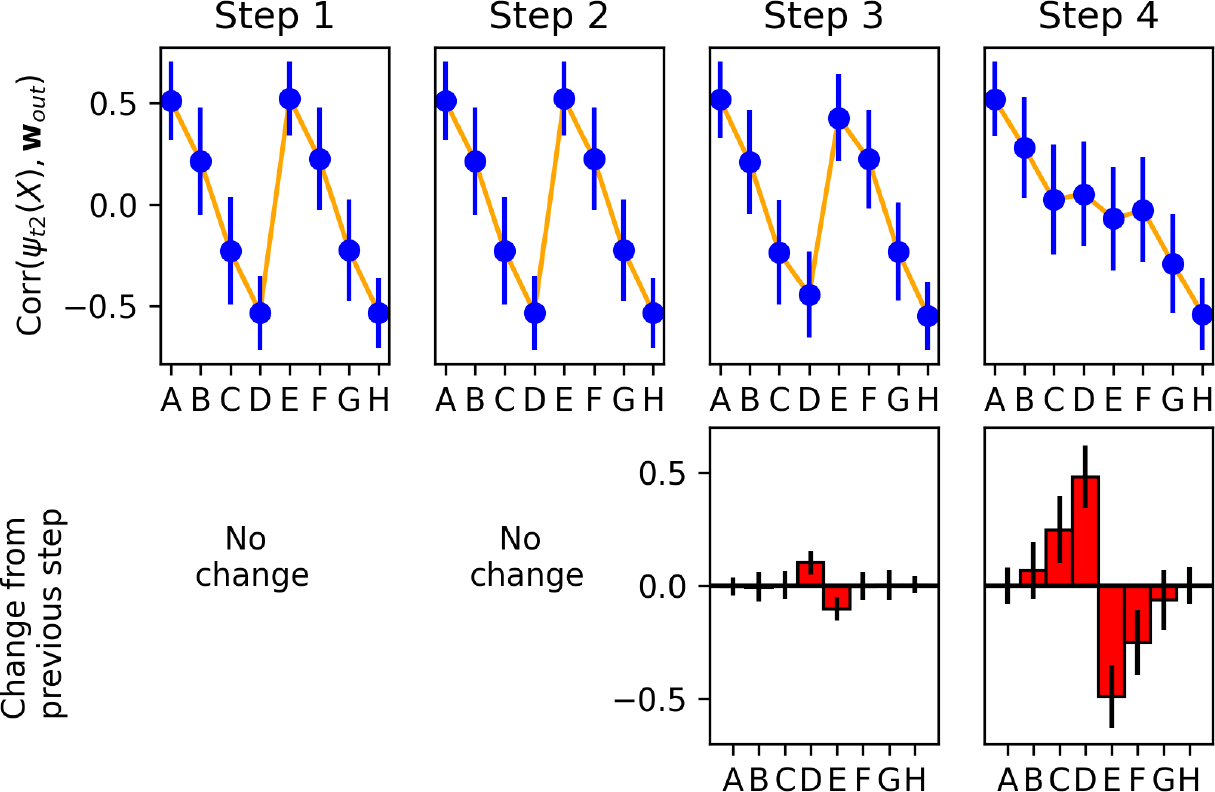
Step-by-step changes in learned representations. Top: Correlation of each item’s step-2 representation ***ψ***_*t*2_(*X*) with the output weight vector **w**_*out*_, measured separately at each time step of trial 20, over 1000 runs in which pair *DE* was shown for the first time on trial 20. Bottom: changes in correlations from one time step to the next (i.e. difference between each upper-row plot and its predecessor). All plots indicate mean and standard deviation over the 1000 runs. Most learning occurs on step 4 (delay step), and learning affects not only the pair shown in this trial (*DE*), but also neighbouring items (*C* and *F*).

By contrast, step 4 produces larger changes, not only for items *D* and *E*, but also for neighbouring items *C* and *F*. These changes are of the correct sign: items *C* and *D* have their representations shifted towards one direction along the decision axis, while items *E* and *F* are shifted in the other direction, as expected.

This shows that information from the currently shown item pair (*DE*) is automatically transferred to other, non-shown items (*C* and *F*). How does the system know which other items should be affected (and in which direction)? A straightforward hypothesis is that the system learns to *couple* adjacent items, with the correct arrangement, as it sees them paired together in previous trials.

To test this hypothesis, we once again look at runs in which the model did not observe item pair *DE* or *ED* before trial 20; but now, out of those runs, we select only the runs in which the model *also* did not see pair *CD* or *DC* before. Fig. S3 shows that for these runs, no transfer to item *C* occurs: at step 4 of trial 20 (when the model first observes pair *DE*), items *D, E* and *F* see some change in their alignment with the decision axis, but item *C* does not. Similarly, not having seen pair *EF* prevents any transfer of information from pair *DE* to item *F*.

These observations confirms that transfer between neighbouring stimuli only occurs if they have been shown together before. Thus, when the model observes a pair of adjacent stimuli, it does not only adjust their learned representations, but also learns to couple them together, so that future trials including either of these items will transfer information to the other (even if this other item is not present in these trials).

### 4.7 Reinstatement of previously paired stimuli in neural activity supports relational learning

The previous results show that the trained network adjusts its learned representations mostly at time step 4, not only for currently shown items, but also for adjacent not-shown items. This raises the question of how this learning actually takes place mechanistically.

The difficulty is that at time step 4, Hebbian learning can only combine information from time steps 2 (as input) and 3 (as output). But at these time steps, the ostensible information encoded in the network is trial response (at step 2) and reward signal (at step 3). None of these contain explicit information about the stimuli being shown (stimuli are only shown at step 1), much less about not-shown neighbouring stimuli. Furthermore, since activations are reset at the start of each trial, no information from previous trials can remain in neural activity. How can Hebbian learning affect representations of not-shown stimuli, as shown in Fig. 8?

We hypothesized that the network must somehow *reinstate* representations of the appropriate items in its neural activity, in parallel to task-related information relevant to producing the correct response for the current trial. That is, neural activations at time step 2 should contain an “echo” of the step-1, feedforward representations ***ψ***_*t*1_ of the items shown at step 1, *and also* of adjacent items not shown at step 1 (all with the appropriate sign). Similarly, step 3 activations should also contain some representation of the output weight vector. This would allow Hebbian learning at step 4 to associate the former with the later, and to stamp this association into the plastic weights (with the correct sign, as specified by *m*(*t* = 4)). This would explain the shift in step-2 representations ***ψ***_*t*2_(*X*), observed at step 4 in Fig. 8. Importantly, this reinstatement would have to occur in parallel with the information ostensibly encoded in neural activations at time steps 2 and 3 (response and feedback signal, respectively).

A difficulty is that the network is not uniformly plastic: each connection has a different baseline nonplastic weight **W**_*i,j*_ and a different plasticity coefficient **A**_*i,j*_. As such, in order to produce adequate learning, the relevant reinstated representations should not be identical to the actual feedforward step-1 input for each item ***ψ***_*t*1_(*X*), or to the output weight vector itself. Rather, they would involve *recoded* versions of these representations, which would produce appropriate learning (i.e. ensure that Hebbian learning pushes step-2 representations of the relevant items in the correct direction along the decision axis), when taking into account the heterogeneous plasticity of each synapse.

Let us call these recoded vectors 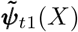 and the recoded version of the output vector 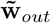 These altered representations can be computed, for each item and for the decision axis, by an optimization procedure (see section F for details). Having found these altered representations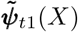 and 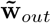 we can simply probe neural activations **r**(*t*) to measure whether any given item *X* is thus represented at any time, by computing the correlation between **r**(*t*) and the altered representation 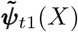

We find that the relevant items are indeed represented in neural activations precisely at time step 2, in their recoded form, and with the appropriate relative signs. Fig. 9 shows the correlation between **r**(*t*) and the recoded step-1 representations 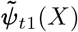 of all possible items *X ∈{A, B, C, D, E, F, G, H,}* for each step of trial 20, using only runs in which the pair shown at trial 20 was either *DE* or *ED*. This confirms that item representations are reinstated in the network’s neural activities (in a suitably recoded state) to support learning for these items. Meanwhile, the recoded representation of the decision axis (output weight vector) is represented specifically at time step 3, with the same sign as the network’s response for this trial (Fig. S4). Together with the error-selective *m*(*t*) signal at *t* = 4, this delayed learning between self-generated, reinstated representations suffices to explain the shift in step-2 representations of these items in the appropriate direction.

**Figure 9.**
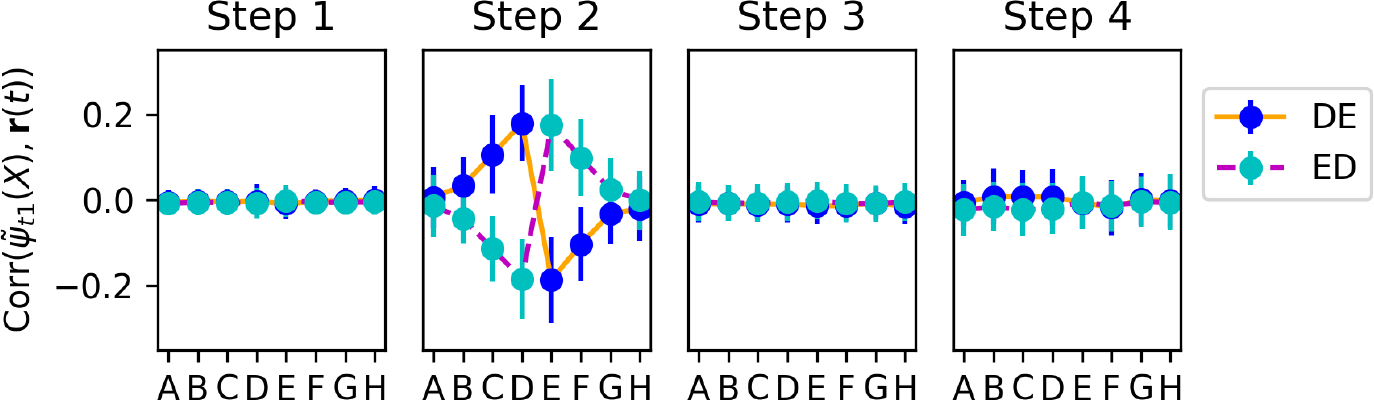
Correlation between **r**(*t*) and 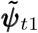 for all items, at each time step of trial 20. Only runs in which trial 20 showed pair *DE* or *ED* are shown (see Fig. 10 for a plot showing all possible pairs) The adapted representations 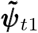 of shown items are reinstated in neural activations at time step *t* = 2, with opposite sign depending on whether the item is item 1 or item 2. Neighbouring items are also reinstated, with appropriate signs.

We stress that these recoded representations 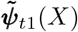 are quite different from the original feedforward step-1 representations ***ψ***_*t*1_(*X*). Correlation between 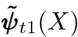 and ***ψ***_*t*1_(*X*) for any item *X* is consistently below 0.1 in magnitude, and can be of either sign. More importantly, if we probe **r**(*t*) for the original ***ψ***_*t*1_(*X*) at each time step, we find no reinstatement at time step 2, or indeed at any step other than step 1 (when the actual items are presented) (Fig. S11). This confirms that the reinstated representations that support delayed step-4 learning are not generally similar to the actual original feedforward representations, though they contain the appropriate information to enable learning for the corresponding items (see section F).

In addition, across the whole batch of 2000 runs, we see a strong relationship between the strength of reinstated, recoded item 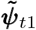 at time step 2, and the magnitude of the shift in step-2 representation ∆***ψ***_*t*2_ that takes place for this same item at step 4 of the same trial (Fig. S12). This confirms that the reinstatement of the item is critical for this item to undergo learning and adaptation of its learned representation.

### 4.8 Coupling of adjacent items for joint reinstatement in future neural activity

This leaves one question: how does the network effect the coupling of adjacent items, so that they can be reinstated together, with the correct sign, in future neural activations? A plausible guess is that the coupling occurs during the presentation of item pairs, which are always adjacent pairs in the early phase of the episode. This seems confirmed by Fig. S3. But how does this take place mechanistically?

We select runs in which the *first* trial in the test episode contained the pair *DE* or *ED*, and again measure the correlation between **r**(*t*) and the adapted vectors 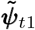 for each item *X*, but this time for trial 1 (Fig. 11. We observe that even at trial 1, when no learning yet has taken place in the plastic weights, step-2 neural activities do reinstate the adapted representations 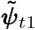 of the two shown items, each with the expected sign. However, only these items are represented: unlike trial 20 (Fig. 9), adjacent items on either side (*C* and *F*) are not reinstated. This shows that, for the items shown in a given trial, step-2 reinstatement is not a learned outcome but an “innate” process, imprinted by gradient descent.

Hebbian learning at step 3 can thus associate the initial feedforward step-1 representations of each item (as provided at step 1) with the reinstated, adapted representation of the *other* item at step 2. As a result, future presentations of each of these items would automatically evoke step-2 reinstatement of the other, now-coupled item, even if it is not present -as observed in Fig.s 9 and 10.

**Figure 10.**
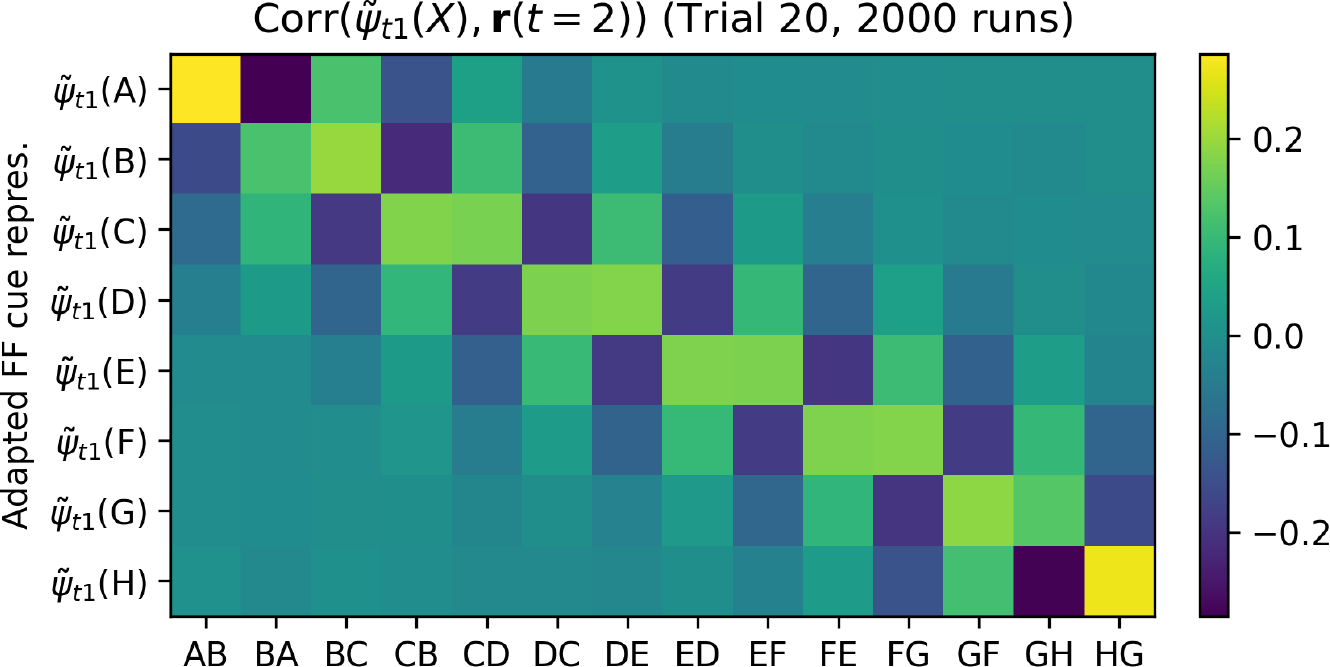
Reinstatement of recoded items, for trials involving all pairs. Correlation between **r**(*t*) and 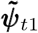 for all items, at time step 2 of trial 20. All runs are shown, grouped separately by which pair was shown in trial 20 for this run. Data for columns ‘*DE*’ and ‘*ED*’ is the same as in Fig. 9 for time step 2.

One difficulty is sign: for this pairing to work as expected, we want the two items to be associated together with the same sign^3^. However, because both items are fed to the network with opposite sign, echoing step-2 adapted representations for either item also have opposite signs (as seen in Fig. 9, 10 and 11). As a result, simple Hebbian learning would couple each item with a step-2 representation of the other item with sign opposite to its own. Whenever one of these items is shown in a future trial, this would lead the adjacent item to be reinstated with the wrong sign. Fortunately, this effect is countered by the fact that *m*(*t*) at step *t* = 3 is consistently negative, independently of reward, as observed in Section 4.2 and shown in Fig. 5. The negative *m*(*t*) inverts the sign of the weight change, and thus ensures that each of the item receives a positive association with the step-2 representation of the other item. As a result, future perception of each item will result in reinstatement of the other with the same sign, ensuring that they receive adjustment in the same direction, as expected.

This also explains the fact that *m*(*t*) at time step 3 (reward time) is largely unaffected by the reward, which might otherwise seem counter-intuitive: step-3 *m*(*t*) does not perform the order learning, but pair coupling between the two items with a view to future joint reinstatement, and thus should have a consistent sign independently of reward for this trial.

**Figure 11.**
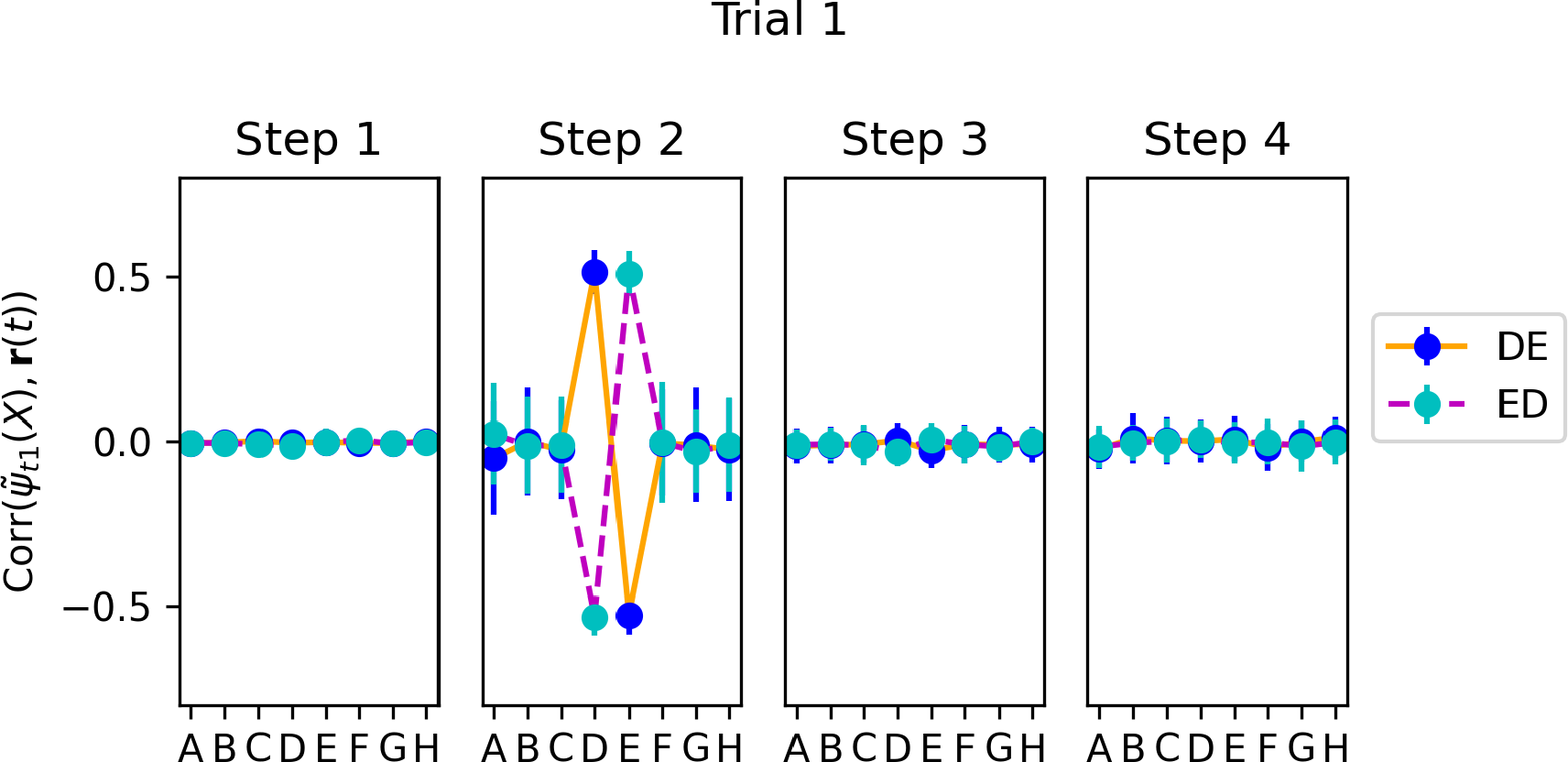
Same as Fig. 9, but for trial 1 (selecting only runs when item pair *DE* or *ED* was shown in trial 1). We see that 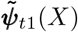 is represented in step-2 neural activity, but only for the two items *D* and *E*.

## 5 An alternative, suboptimal solution cannot perform list-linking

As shown in Fig. 3, the optimization process sometimes settle on a less efficient solution. This alternative solution turns out to be similar to the simple, passive reward-modulated Hebbian learning process suggested in section 4.6.1: *m*(*t*) is strongly sensitive to reward at *t* = 3 (reward delivery time) and modulates Hebbian learning between feedforward item input (at step 1) and response (at step 2) (see Fig. S7). Representation learning only seems to occur at time step 3 (reward delivery time) (Fig. S9). Thus, this solution dispenses with active reinstatement of previous representations, relying only on present-trial information for learning.

Importantly, this “passive”, or reactive solution fails on the list-linking task, unlike the active, cognitive solution described previously. Because the passive solution only involves present-trial information, it is unable to quickly reassemble existing knowledge upon learning new information that does not explicitly reference this existing knowledge. See Appendix H for details.

Across multiple experiments and parameters, we found that the training process consistently produces one of these two discrete outcomes (high-performance cognitive solution based on reinstatements vs. less accurate reactive solution based on reward-modulated learning). Nearly all trained network settled on one of these two solutions.

## 6. Discussion

### 6.1 A discovered cognitive mechanism for relational learning and fast knowledge reassembly

How does the brain learn underlying relationships and thus gain insight from limited experience? In this study, we investigated two classical relational tasks — transitive inference and list-linking — for which there are few if any known biologically realizable mechanisms. To discover such mechanisms, we took a meta-learning approach in a neural network architecture that captures the brain’s ability to control its own learning across task trials. Because we had full access to the network’s internals, we were able to obtain a complete mechanistic understanding of how the trained network learns and represent relations. Our main findings were as follows:

- The trained networks successfully learn orderings from pairwise presentations and perform transitive inference, exhibiting the symbolic distance effect and serial position effect commonly reported in behavioral experiments.
- Through learning, stimuli are mapped to learned vectors, whose alignment with the output weight vector encodes item rank. Pairs are represented by subtracting these learned representations of individual items, automatically encoding difference in rank.
- In the dominant, high-performance solution, this learning involves a cognitive act: previously seen items are actively and selectively reinstated in working memory (in recoded form), in parallel with task execution, supporting delayed representational learning.
- This delayed, self-generated learning allows the network to adjust representations not just for currently shown stimuli, but also for other items shown in the past, supporting fast reassembly of existing knowledge.
- An alternative solution based on passive, reactive learning (reward-modulated Hebbian association of external stimuli) can perform transitive inference (though at a lower level of performance), but cannot reassemble existing knowledge about stimuli other than those currently shown.

The network uses selective reinstatement of previous items to support delayed learning and knowledge assembly. Such reinstatement is reminiscent of neural activity that appears to reflect re-activation of memory traces, most notably “replay” in neural firing and similar activity patterns seen in the hippocampus [Buzsáki, 2015, Foster, 2017]. Importantly, these activity patterns have been proposed to subserve not only consolidation in memory of specific past experiences [O’Neill et al., 2010, Carr et al., 2011, Ó lafsdó ttir et al., 2018], but also construction of abstract internal representations enabling inference and generalization [Buckner, 2010, Buzsáki, 2015, Joo and Frank, 2018, Comrie et al., 2022, Kurth-Nelson et al., 2023]. A notable difference is that, in the present model, reinstatement occurs predictably at fixed time points in a task trial, whereas replay in the brain occurs spontaneously, and indeed even outside of periods of task engagement. Nonetheless, since reinstatement is self-generated, it constitutes a formal bridge to internally generated neural activity patterns which may have a role in memory and generalization [Buzsaki, 2006, György Buzsáki, 2019].

### 6.2 Implications for experiments

The mechanism elucidated above may suggest new interpretations and analyses of neural data. The model suggests that transitive inference benefits from the reinstatement of other items than those presented in the current trial. Such a reinstatement process may be investigated in neural data. However, importantly, our results point out that these reinstated representations need not be identical, or even similar, to those directly elicited by stimulus presentation. Because the function of these reinstated representations is to support learning (and limit interference with ongoing task performance), rather than recapitulate sensory-driven input, they may be *recoded* in a way that greatly differs from the sensory representations of these same items (compare Fig. 9 and S11).

Here we hypothesized that the items would be recoded in a form that supports learning in the face of heterogeneous synaptic plasticity, and derived an optimization-based procedure to find these recoded representations. While our approach successfully identified reinstated items, this recoding is also likely bound by additional constraints. One such constraint is to minimize interference with the ongoing main task, i.e. choosing between the two stimuli in the current trial (see Libby and Buschman [2021] for an observed example of sensory representations being recoded to orthogonal memory representations in a working memory task). Thus, different approaches may prove more suitable for probing experimental neural data (see Section F).

More generally, the discovered solution exemplifies the importance of geometrical, whole-population approaches. The mechanism relies on manipulating vectors in neural population space, which could not be identifiable from purely single-cell analyses.

### 6.3 Meta-trained plastic neural networks as models of cognitive learning

Finally, our results illustrate the potential benefits of modelling approaches based on *plastic* artificial neural networks. Many recent studies have adopted training recurrent neural networks (with external, hand-designed algorithms) to identify rich dynamics capable of performing various tasks [Mante et al., 2013, Yang et al., 2019, Duncker et al., 2020]. The present paper extends this approach to plastic networks, capable of controlling their own learning, allowing gradient descent to discover not just representations and dynamics, but also the learning processes that acquire these representations and dynamics. These discovered learning processes can then be fully elucidated by judicious experimentation on the trained networks. The results reported in this paper demonstrate the power of this approach, and suggest that training plastic neural networks offers a potentially fruitful avenue of investigation into the space of neural mechanisms that support learning and cognition.

## Acknowledgements

We thank Larry Abbott, Vincent P. Ferrera and Greg Jensen for fruitful discussions about this work and comments on an earlier version of the paper.

This work was supported by the the Simons Collaboration for the Global Brain (521921 and 542981), the NSF NeuroNex award (DBI-1707398), the Gatsby Charitable Foundation, and an NIMH K99 (MH126158-01A1, KK).

## Supplementary Materials

### A Learning multiple lists

We ensure that the networks can learn multiple orderings rather than just one. This is done with two interventions. First, we only reset plastic weights **P**(*t*) every third episode. Second, the “test” period of each episode (the last 10 trials) include 25% of pairs involving stimuli from the previous ordering since the last **P**(*t*) reset (if any). While not immediately relevant to the present work, this constitutes a simple example of meta-continual learning, where the goal is not just to be able to learn new problems efficiently, but also to keep in memory previously seen problems, at least up to a short horizon. Performance on pairs from previous lists was consistently high (above 70%), confirming that the model is able to learn and store several different lists.

The above procedure was followed for all training. At test time, among the results reported in this paper, the list-linking examples are implemented as three successive episodes of different lengths (plus a single test trial), without initialization of **P**(*t*) between these episodes. All other reported results used a single test episode with **P**(*t*) initialized to 0 at the start of the episode.

### B Meta-Reinforcement Learning algorithm: The A2C algorithm

The outer loop of our method ascends the gradient of episodic reward over the network’s structural parameters (the various **W** matrices and **A**, plus additional scalar parameters; see below). Gradients are estimated by A2C (Advantage Actor-Critic), a well known reinforcement learning (RL) algorithm that is commonly used in meta-RL experiments [Duan et al., 2016, Wang et al., 2016, Miconi et al., 2018, Wang et al., 2018]. First, in addition to previously discussed outputs, the network also produces a scalar output 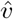 (the “value-prediction” output), through the weight vector 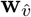. This output tries to predict, at each point in time, the current value of 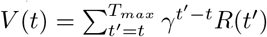 that is, the sum of future discounted rewards during the episode. After the episode concludes, we retroactively go back through the history of rewards obtained at each time step, allowing us to compute the real, ground-truth *V* (*t*) for each time step. Then the actual objective *J* to be maximized is computed as follows:

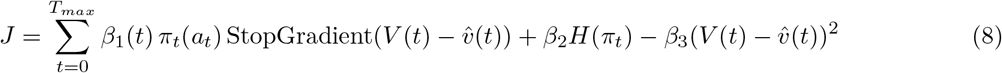

Here, *a*_*t*_ is the action (i.e. the response) given at time *t, π*_*t*_ is the probability distribution over both possible actions/responses (determined at each time step by a SoftMax over both output neurons of the network, see Methods). *H* denotes a measure of dispersion of a probability distribution, usually implemented as the Shannon entropy; in practice, due to computational limitations, we set the *H* term as the sum of squares (an alternative measure of dispersion) rather than the actual Shannon entropy. *β*_1_(*t*) is a multiplier set to 1 for the first 20 trials (on adjacent pairs only) and 4 for the last 10 trials (on all pairs; see Methods), and *β*_2_, *β*_3_ are positive constants. Note that the second term encourages the output distribution over possible actions to be as uniform as possible (with the goal of favouring exploration), while the third term optimizes the value predictor head 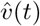

Intuitively, this algorithm can be thought of as a version of the standard REINFORCE method for estimating the gradient of rewards over weights that control stochastic responses: take the weight modification that would make the response you just gave more likely in the future for the same inputs, and multiply it by the actual rewards obtained. Furthermore, in A2C (as in other actor-critic methods), the actual rewards are centered by subtracting a network-generated estimate of future discounted rewards 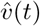 This centering has the effect of reducing the variance in the gradient estimate. In addition to this, training the value predictor also plays the role of an “auxiliary loss”: a signal with a hard ground truth that provides information about environment structure. Finally, A2C also includes an additional entropy term *H*(*t*) to encourage maximally uniform distribution over possible outputs, and thus (hopefully) exploration.

The parameters being subject to gradient descent between episodes are **W** matrices, **A**, *η*, and a scalar multiplier on *m*(*t*). Gradients of *J* over these parameters are computed automatically by PyTorch, using the Adam optimizer. Note that gradient computation involves backpropagation of gradients through time, since activities at time *t* influence activities at future times. One step of gradient descent is applied after each episode, backpropagating through time over all 30 trials of the episode. We reiterate that no gradient descent occurs during a given episode.

**Figure S1:**
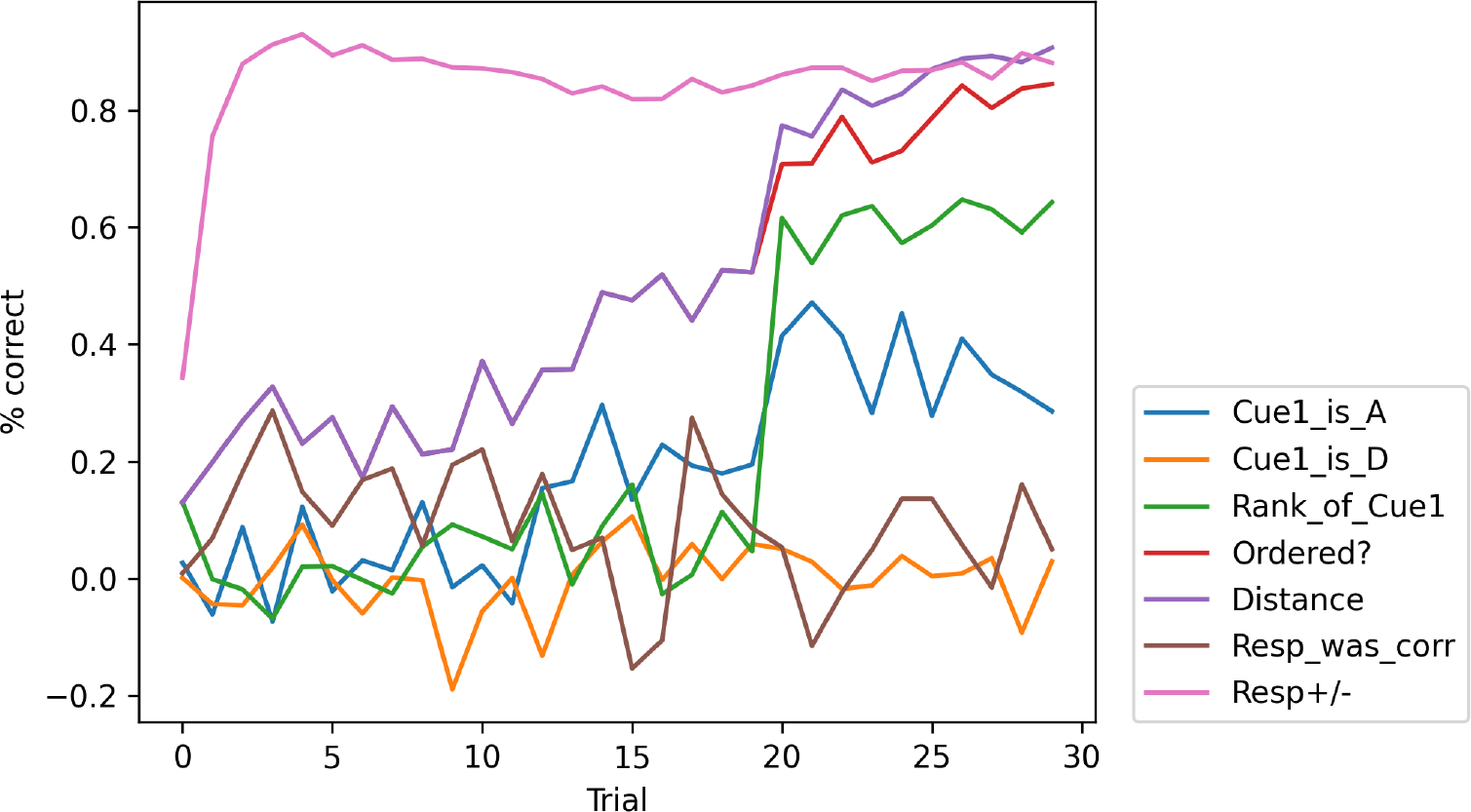
Decoding performance, at each trial in the episode, for various quantities. See text for details.

### c. Individual item rank is not directly encoded in neural representation of pairs

We run a full episode with a trained network over 2000 runs (each with different randomly generated stimuli), and use a linear decoder to decode various quantities from **r**(*t* = 2) (step 2 neural activations), separately for each trial. We train our decoder on data from the first 1800 runs, and test it on data from the last 200 runs.

Results are shown in Fig. S1. The quantities we try to decode are:

- *Cue1_is_A*: whether item 1 in this trial is *A*.
- *Cue1_is_D* : whether item 1 in this trial is *D*
- *Rank_of_Cue1* : the actual numerical rank of item 1
- *Ordered?* whether the items were shown in descending order (Stim1 ¿ Stim2) in this trial
- *Distance*: the distance in rank between the items (which is always 1 for the first 20 trials)
- *Resp_was_corr* : whether the network’s response was correct for this trial
- *Resp+/-*: the network’s response for this trial

Information about the network’s response is clearly encoded in **r**(*t* = 2) from early on in the episode (pink curve). Information about the ordering of the two items (i.e. the actual correct response for this trial) and their distance slowly increases over time (purple and red items). Note that before trial 20, all pairs shown are adjacent pairs, and thus item ordering and item distance (always −1 or +1) carry the same information, explaining their identical curves up to this point.

By contrast, information about the identity of each individual item is not encoded at all before trial 20. Even after trial 20, the apparent ability to decode item 1 rank is very similar to that of a classifier only given the correct response for this trial as input; that is, **r**(*t*) does not seem to possess additional information about individual item rank above and beyond that provided by knowledge of the correct response for this trial (data not shown). Thus, the network does not seem to reliably encode individual item identity in its responses.

### D Representation changes for item pairs other than *DE*

In Fig. S2, we show step-by-step representation changes at trial 20 when trial 20 involved the pair *CD* (top) or *EF* (bottom), rather than *DE* as in Fig. 8. We see that results are very similar: representation changes occur at time step 4, and involve not just the items shown at this trial, but also adjacent items (*B* and *E*, or *D* and *G*, respectively).

### E Transfer of representation changes to unseen neighbouring items requires previous joint presentation

In Fig. S3 (**a**), we plot the same ste-by-step changes in representation as in Fig. 8. However, instead of including all runs in which the model did not observe item pair *DE* or *ED* before trial 20, we now only include runs in which the model observed neither *DE/ED* nor *CD/DC* (item pair shown at trial 20 is still *DE* or *ED*). The figure shows that for those runs, no transfer to item *C* occurs: at step 4 of trial 20 (when the model first observes pair *DE/ED*), items *D, E* and *F* see some change in their alignment with the decision axis, but item *C* does not.

Fig. S3 (**b**) shows the same experiment but now the additional withheld pair is *EF* instead of *CD*. Now learning does not transfer to item *F*, as expected. This confirms that transfer between neighbouring items only occurs if they have been shown together before.

**Figure S2:**
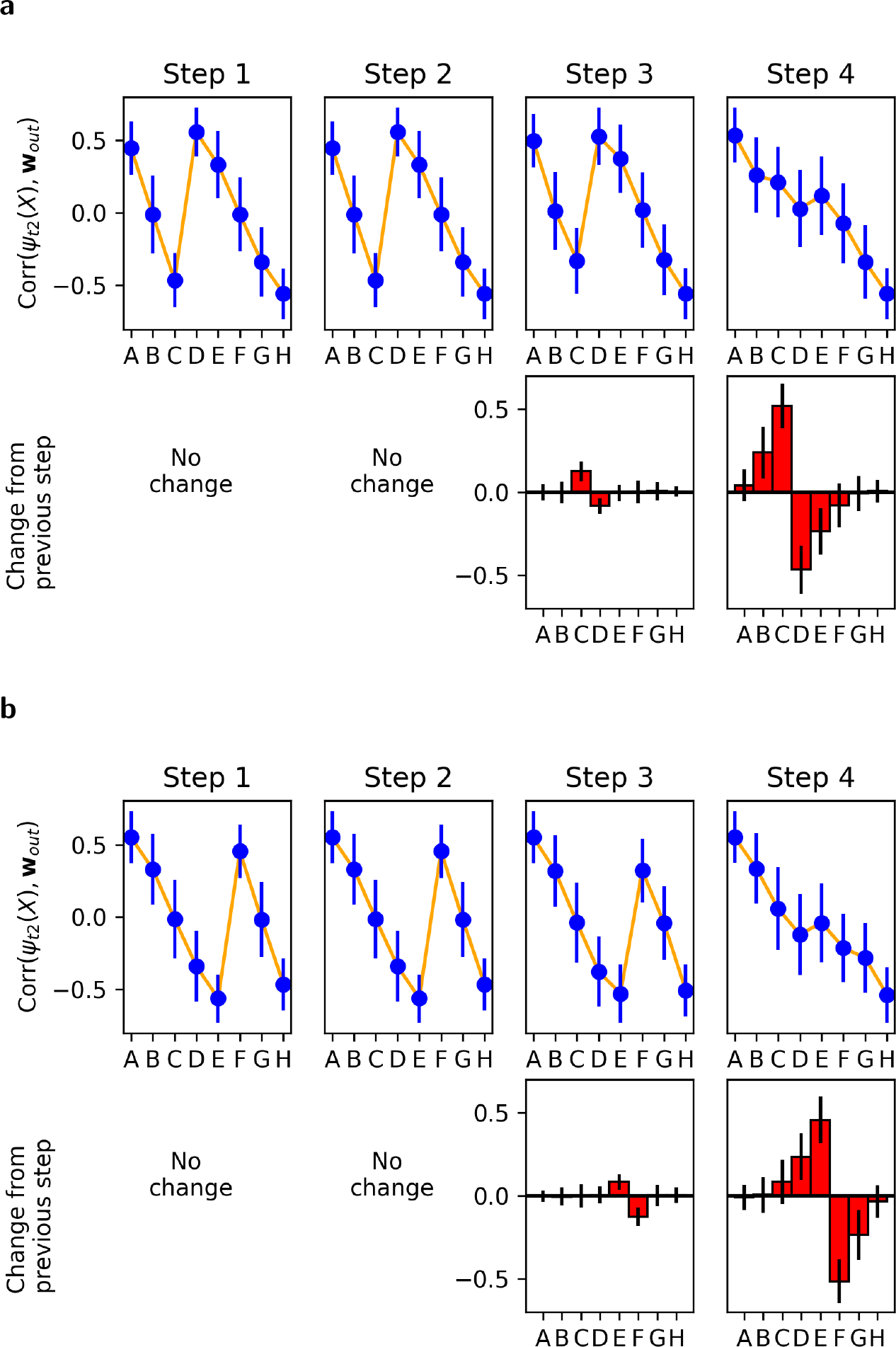
Step-by-step changes in learned representations at trial 20, for different withheld pairs. Conventions are as in Fig. 8, but here the pair that is withheld until trial 20 is either *CD* (**a**) or *EF* (**b**) rather than *DE*. Learning proceeds similarly for these other pairs.

**Figure S3:**
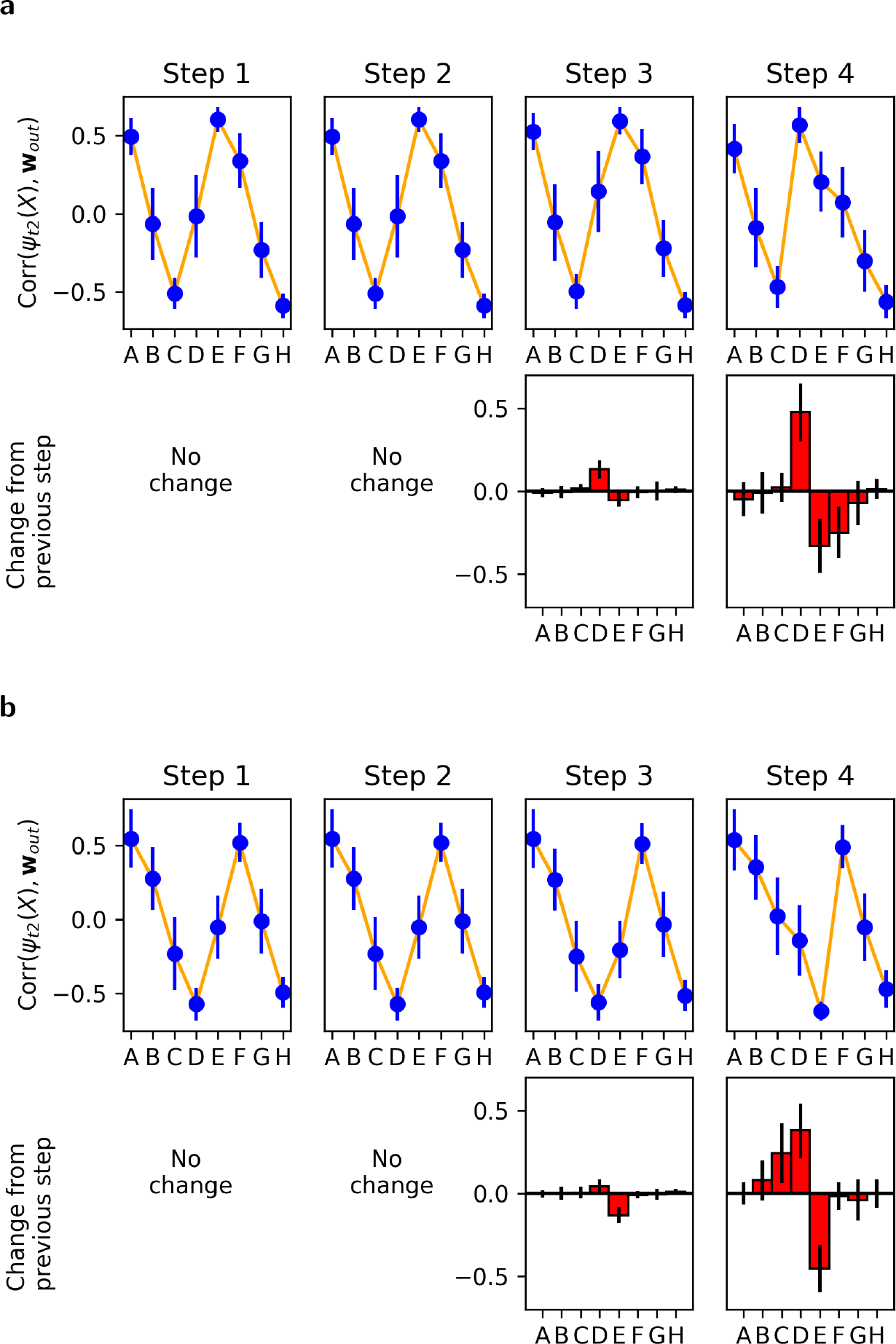
Step-by-step changes in alignment when showing pair *DE* (in any order) at trial 20, after withholding both pair *DE* as well as a neighbouring pair until trial 20. **a**: additional withheld pair is *CD*. **b**: additional withheld pair is *EF*. At time step 4, and unlike Fig. 8, no transfer of learning to the withheld neighbouring item occurs. This confirms that transfer of information from currently shown items to neighbouring items only occurs if these other items have been shown in pair with the currently shown items before.

**Figure S4:**
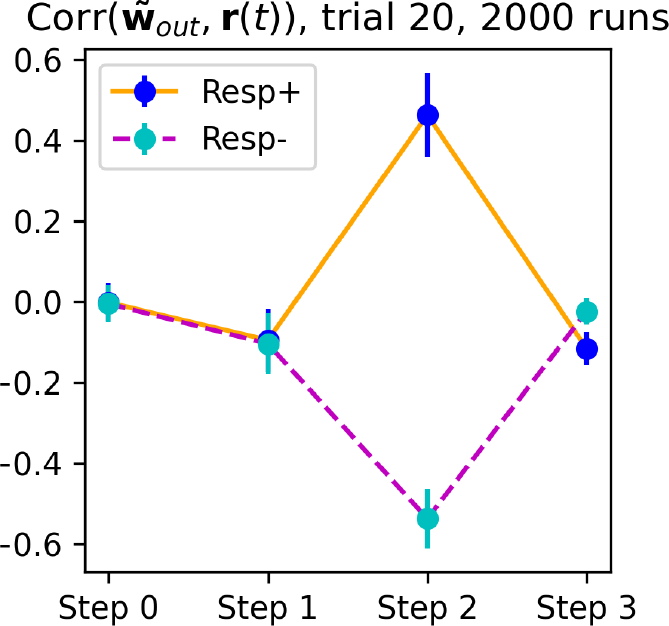
Correlation between **r**(*t*) and 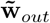 at each time step of trial 20 (2000 runs), shown separately for runs in which network response at trial 20 was positive vs. negative. The adapted representation of the output weight vector 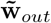 is strongly represented at step 3, with sign correlated with network response for this trial.

### F. Identifying recoded representations with an optimization procedure

Our analysis indicated that representation learning occurred at time step 4, and involved a shift in the step-2 representation of relevant items along the axis of the output weight vector (with either sign) (Fig. 8). These step-2 representations, in turn, are the result of passing step-1 feedforward representation through one step of recurrence with the learned plastic weights **P**(*t*). Thus, if the step-2 representations change, it implies that **P**(*t*) also changed. How did this change happen?

Recall that plastic weights are shaped by Hebbian learning, which essentially amounts to **∆P**(*t*) *≈ m*(*t*) **x**(*t−* 1) **r**(*t−* 2)^T^ (see Equation 7). For intuitive understanding, let us overlook the nonlinearity and assume a positive *m*(*t* = 4). If weight changes were purely Hebbian, including the step-1 representations in **r**(*t* = 2) and the output weight vector in **r**(*t* = 3) would increase plastic weights at time *t* = 4 by 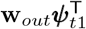 which would have exactly the appropriate effect for representation learning: future presentations of 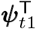 to the recurrent network (at *t* = 1)would produce an output (**r**(*t* = 2)) more similar to **w**_*out*_ (vs. less similar for a negative *m*(*t*)). This would constitute a straightforward example of Hebbian associative learning.

However, as seen in Fig. S11, this does not occur. Neural activities at time steps 2 and 3 do not seem to contain step-1 representations of the items, or of the output weight vector. This leads to the following question: what signal in the neural activities at time step 2 and 3 produces the representation changes seen at step 4 in Fig. 8 ?

We hypothesized that the network does reinstate representations of the items and decision axis, but under a *recoded* form, which would facilitate learning under the constraints imposed upon the network. One such constraint (we surmised) is heterogeneous plasticity. Recall that the recurrent weights are not purely Hebbian, but are rather the sum of a fixed (non-plastic) **W**_*rec*_ and a weighted plastic component **A** *⊙* **P**(*t*) (see Equation 1). Thus, simple Hebbian learning between ***ψ***_*t*1_ at time *t −* 2 and **w**_*out*_ at time *t −* 1 would indeed shift the **P**(*t*) component in the correct direction (towards increasing alignment between ***ψ***_*t*2_ and **w**_*out*_), but the net effect on the *total* recurrent weights **W**_*rec*_ + **A** *⊙* **P**(*t*) would be more complex and unpredictable. To take an extreme example, if **A** were fully-negative and larger than **W**_*rec*_, the actual effect on total weights would be in the opposite direction.

What would be the “ideal” activation vectors **r**(*t−* 2) and **r**(*t −*1) that would produce the desired weight change in *total* recurrent weights at time step *t*? Since no obvious analytical answer is available, we address this question by an optimization procedure. Recall that the desired change is to shift future step-2 representations of each item towards greater alignment with the output weight vector. Therefore, we seek to find the recoded representations 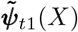 and 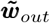 which, when represented in network activity at times *t −* 2 and *t −* 1 (and thus modifying **P**(*t*) proportionally to their outer product, due to Hebbian learning) would cause a net change in the *total* recurrent weights, such that future presentations of the original ***ψ***_*t*1_(*X*) would produce an output more similar to the original ***w***_*out*_.

We initialize 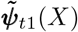 (for all items *X*) and 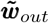 to small random vectors, and repeatedly perform the following procedure:

1. Compute the outer product 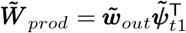
2. Substituting this outer product for the plastic weights **P**(*t*) in the total recurrent weights, recompute the recurrent operation on the original step-1, feedforward representations: 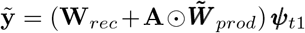 (recall that **W**_*rec*_ and **A** are fixed within each episode)
3. Compute the alignment between this output and the decision axis 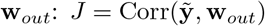
4. Perform one step of gradient ascent of *J* over 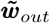 and 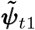

This procedure is performed in parallel over the batch of 2000 individuals, each with randomly generated stimuli. Because each run has its separate stimuli, we maintain a separate set of 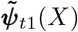 for each of the 2000 runs. By contrast, we consider 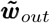 as an intrinsic feature of the network, and thus use only one 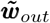 for the whole batch. We use the Adam optimizer with learning rate 1e-3 and weight decay 1e-2 for regularization. It is important to note that these recoded representations were not found by specifically exploring network activity at time steps 2 and 3. Rather, they constitute hypotheses about what learning-appropriate representations at these times “should” look like, at least in part. Clearly many other factors affect the structure of ideal representations, e.g. not interfering with the main task: the reinstated representations should minimize interference with ongoing task-relevant signals, i.e. response at time step 2 and reward at time step 3 (see Libby and Buschman [2021] and references therein for examples of interference-minimizing recoding of representations in neural data). From Fig.s 9, 10, and S4, we get confirmation that our “guessed” representations, based on heterogenous plasticity alone, are adequate probes to detect the reinstatement of items by the network at these time steps, whereas the original unadapted ***ψ***_*t*1_(*X*) and ***w***_*out*_ are not (Fig. S11).

### G. Robustness to massed presentation

Some existing models of transitive inference are disrupted when one single pair is shown a large number of times [Jensen et al., 2019]. Following existing practice, we ran the trained network on an extended episode, in which 20 additional trials (all with the pair *DE*) were inserted between the 20 adjacent-only trials and the 10 all-pair trials. No other modifications were made. Again, we performed 2000 runs, each with independently generated stimuli. As shown in Fig. S5, there was little impact on performance, whether on transitive inference, the symbolic distance effect, or the end-item effect.

**Figure S5:**
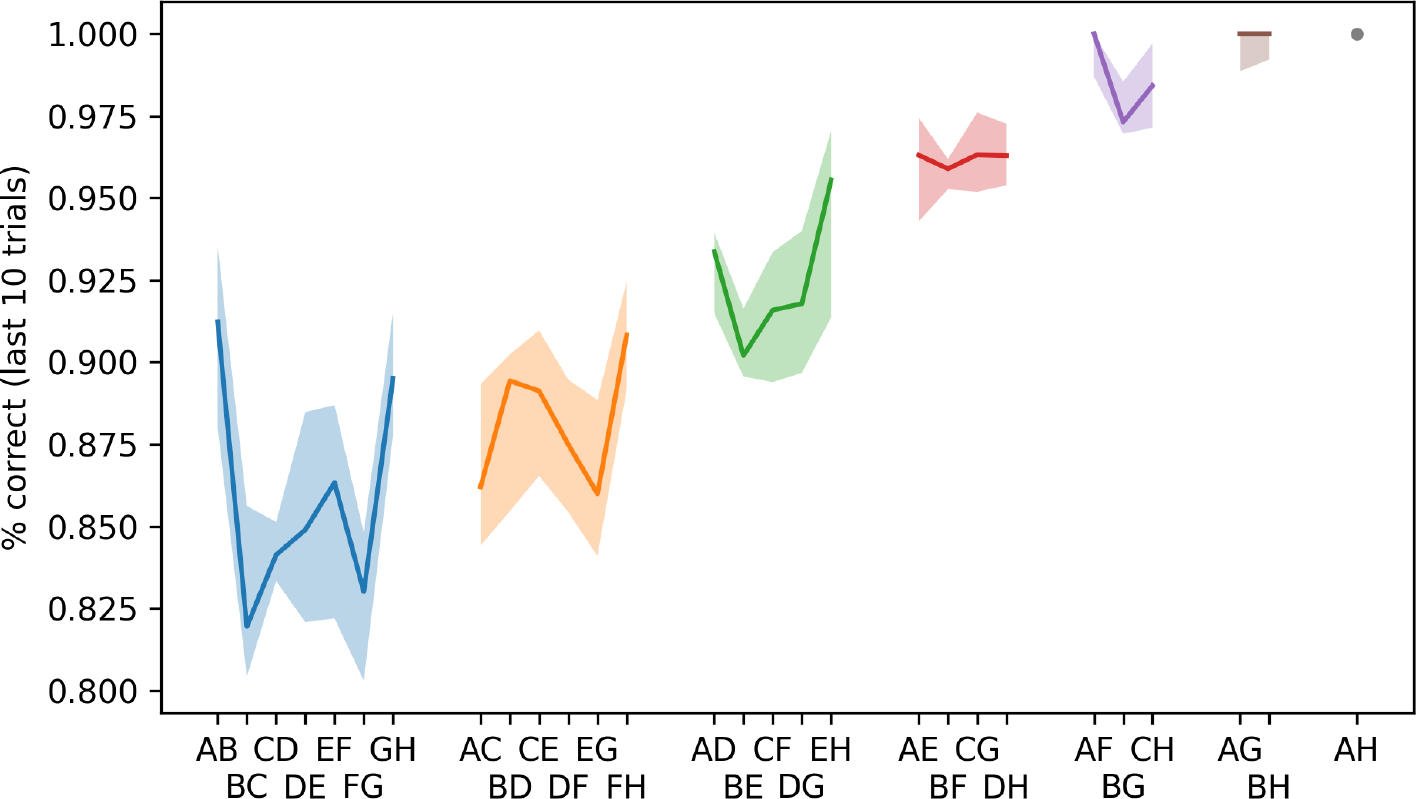
Performance for massed-presentation experiment, grouped by pair. Compare with Fig. 3.

**Figure S6:**
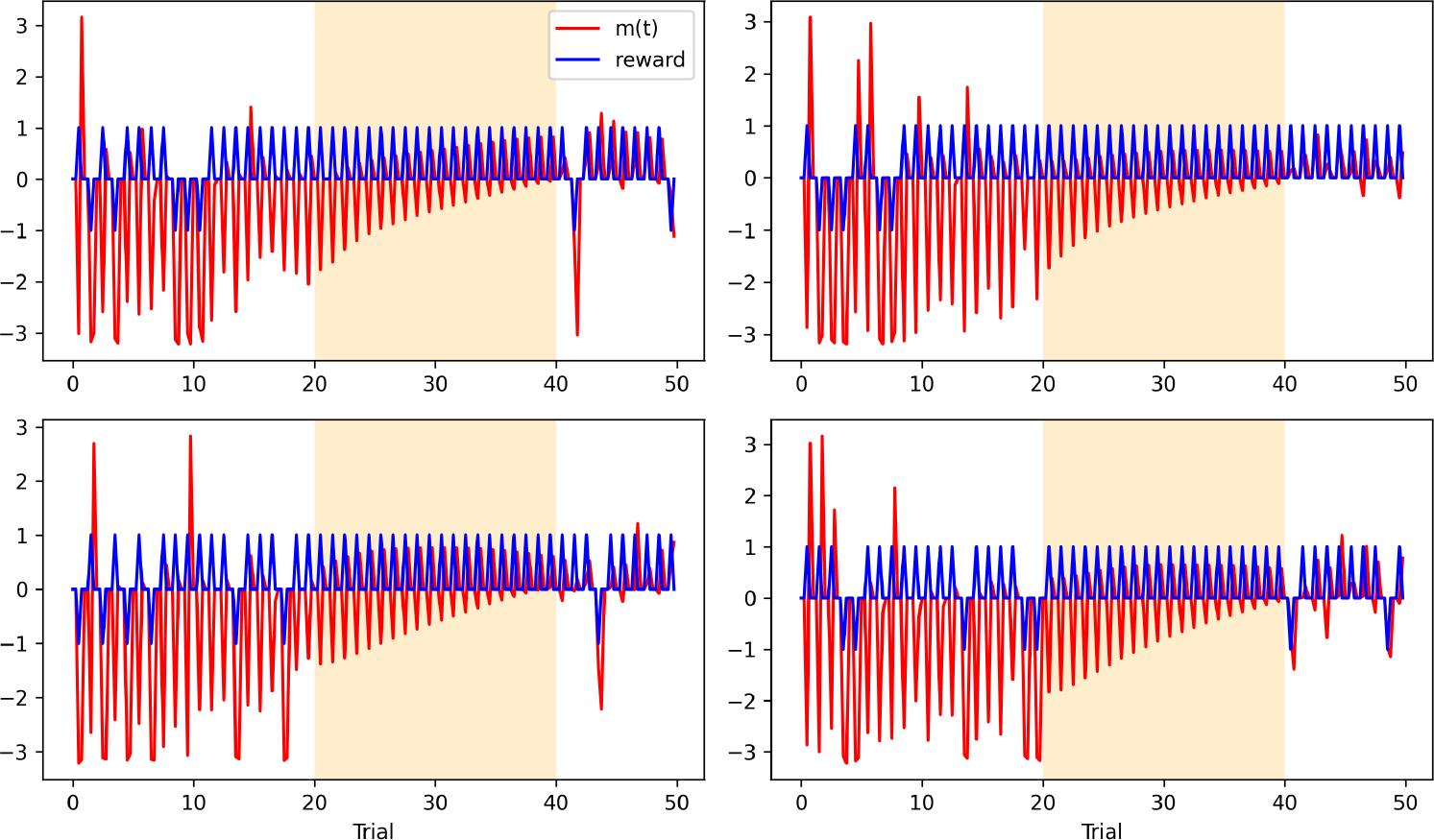
Traces of neuromodulatory output *m*(*t*) (red) and reward signal (blue) for 4 episodes with massed presentation of the pair *DE* between trials 20 and 40.

**Figure S7:**
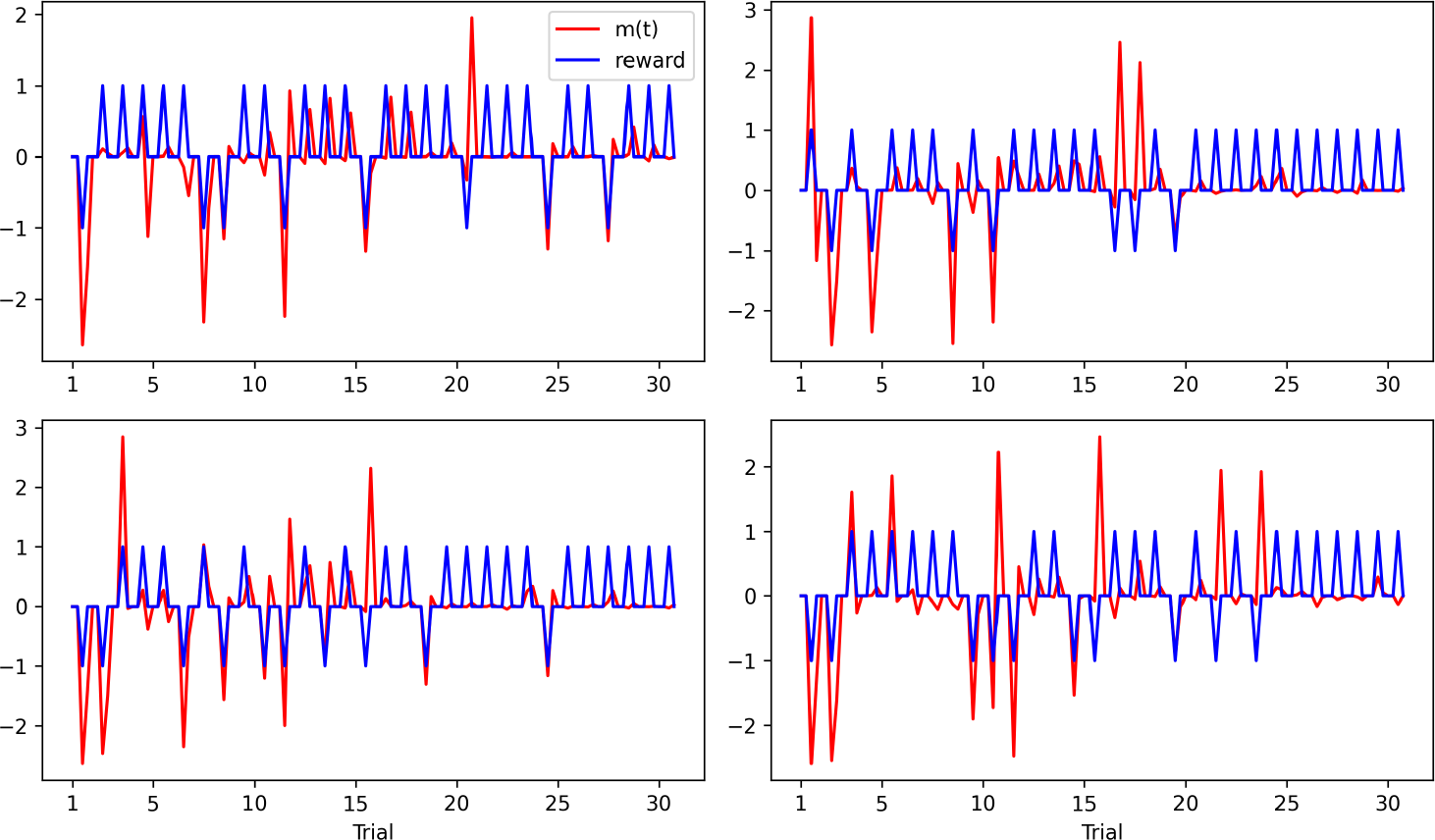
Traces of *m*(*t*) (red) and reward signal (blue) over one episode, for four different runs, using a suboptimal network. Conventions are as in Fig. 5

### H. An alternative, suboptimal solution

Here we describe results from the alternative, suboptimal solution sometimes found by the system, as discussed in section 5. As shown in Fig. S7, this solution uses reward-selective neuromodulation *m*(*t*) at reward time, while *m*(*t* = 4) seems unrelated to reward.

Fig. 7 shows that this alternative solution still maps each single item to a learned representation that encodes its rank, though with a somewhat flatter curve near the middle of the series. As shown in Fig. S10, the network still exhibits a symbolic distance effect, confirming that the overall representational scheme is similar. However, performance in list-linking conditions is at or below chance.

What might cause this deficit in list-linking? In Fig. S9, we see that representation changes at a given trial occur at reward time only. Importantly, representations are only shifted for the items presented at the current trial. This explains failure of list-linking (which requires transfer of information from currently shown items to other not-shown items).

As mentioned in the main text, almost all meta-trained networks followed one of the two solutions described in the present paper, up to mutually compensating sign changes that leave the fundamental mechanism invariant. The balance of the two solutions may change according to various experimental choices (for example, forcing **A** to contain only positive values increased the proportion of suboptimal solutions). As shown in Fig. 3A, the process as described inthe Methods greatly favours the higher-performance cognitive solution over the passive one.

**Figure S8:**
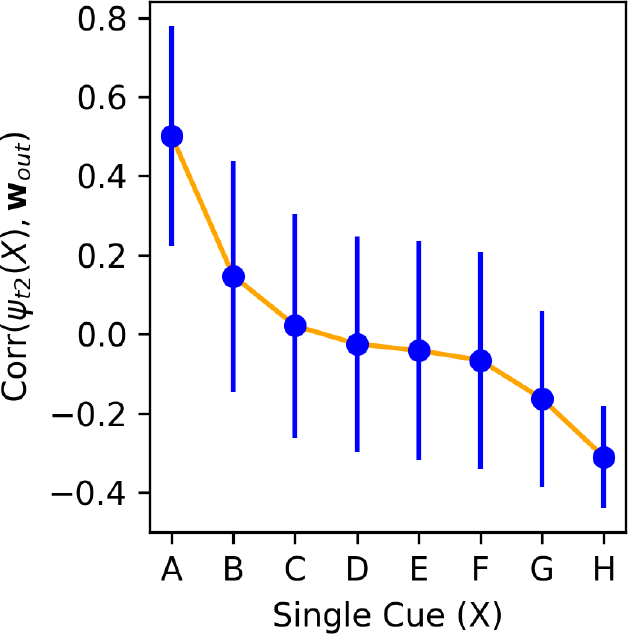
Correlation between the step-2 representation of each item ***ψ***_*t*2_(*X*) and the output weight vector **w**_*out*_, at trial 20, for the suboptimal network. Alignment between step-2 representation and output weight vector still monotonically encodes rank, but with a flatter profile than in the standard network (see Fig. 7).

**Figure S9:**
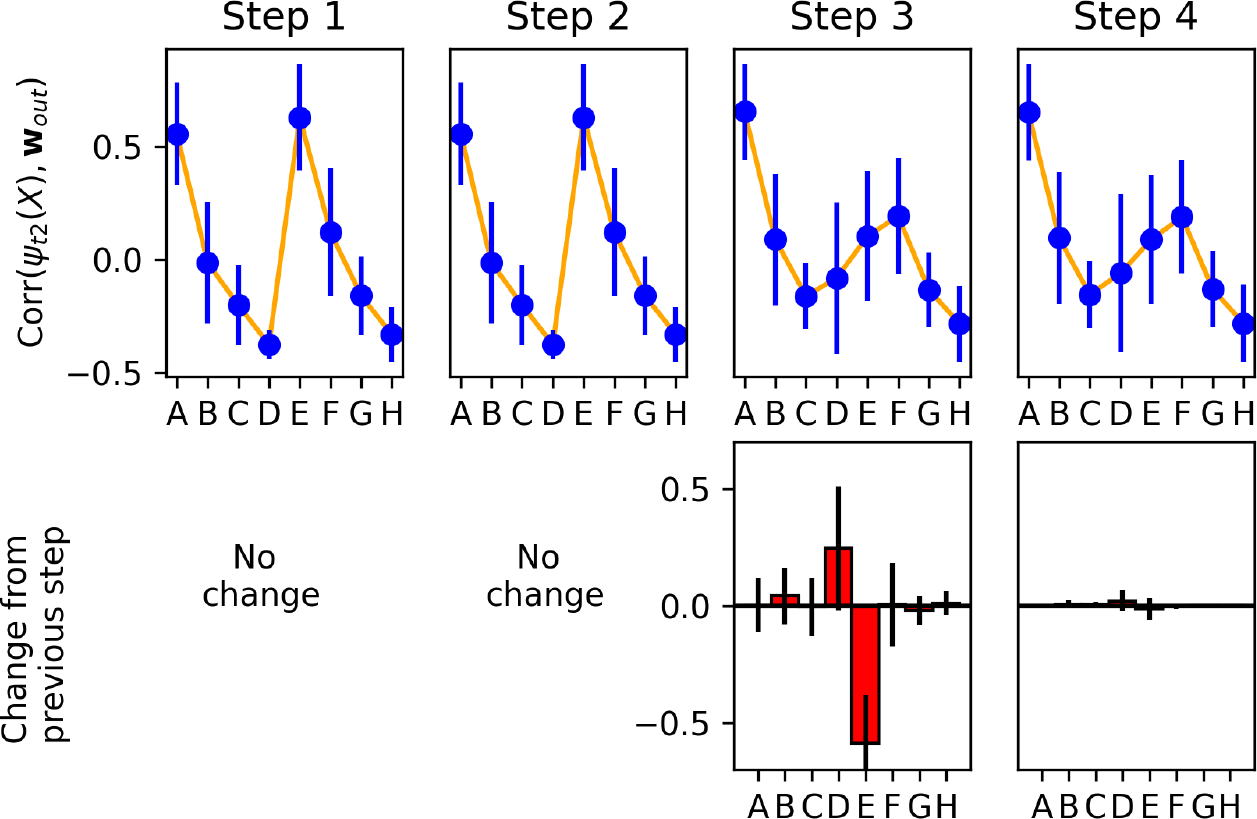
Same as Fig. 8, but for the suboptimal network. Representation changes occur at time step 3 (reward delivery) rather than time step 4, and only for the items shown in the current trial.

**Figure S10:**
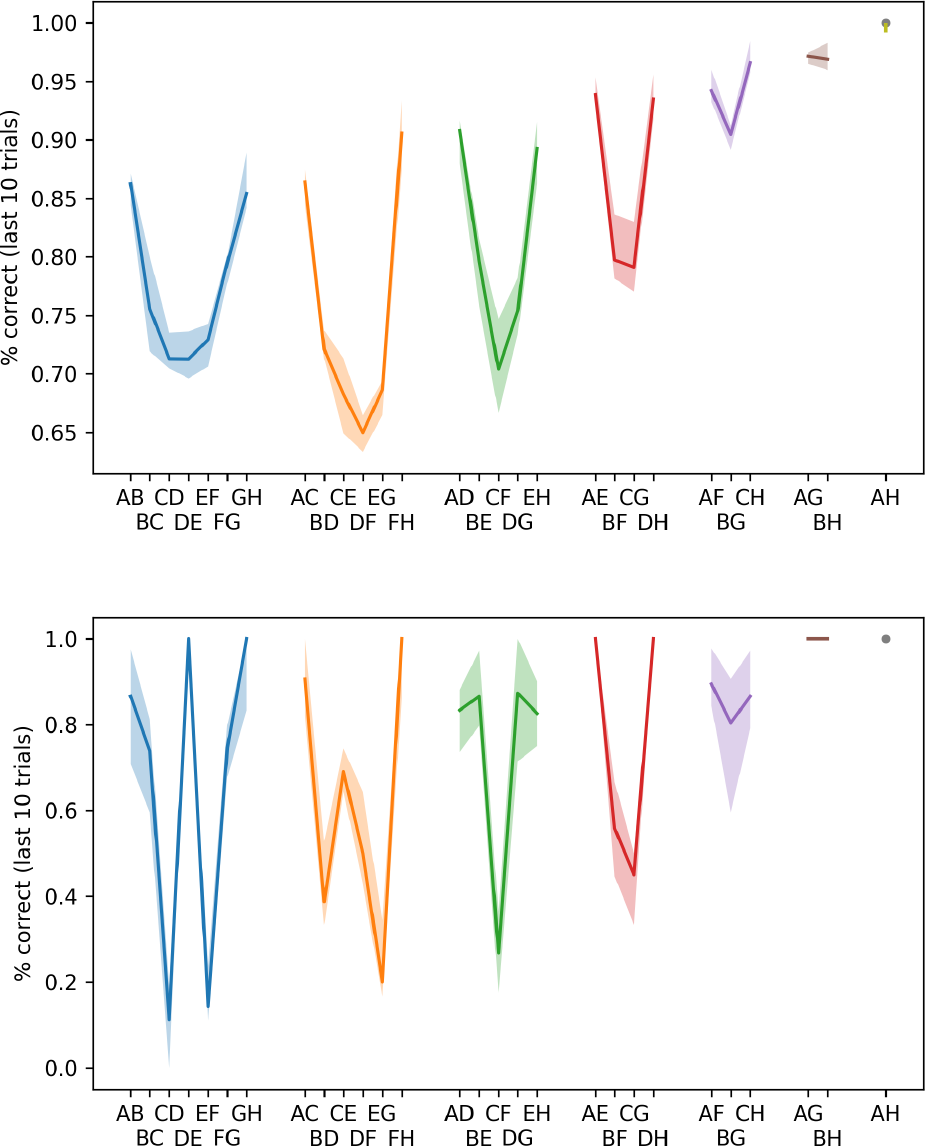
Performance at test time grouped by pairs, in normal conditions(top) and list-linking conditions (bottom). The suboptimal network is still able to produce a symbolic distance effect in normal conditions, but fails the list-linking task. Conventions as in Fig. 3 and 4.

**Figure S11:**
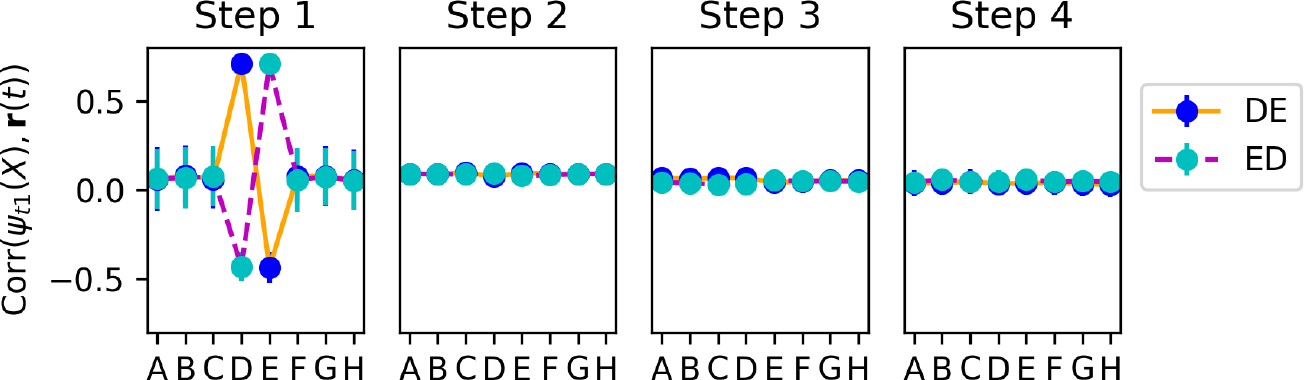
Same as Fig. 9, but using *ψ*_*t*1_ (the original step-1 feedforward representations of each item) instead of 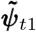 (the adapted heterogeneity-aware representations, found by optimization). Again, only runs in which trial 20 showed pair *DE* or *ED* are shown. Unadapted, original step-1 representations are strongly present at step 1 (unsurprisingly), but not at any other step.

**Figure S12:**
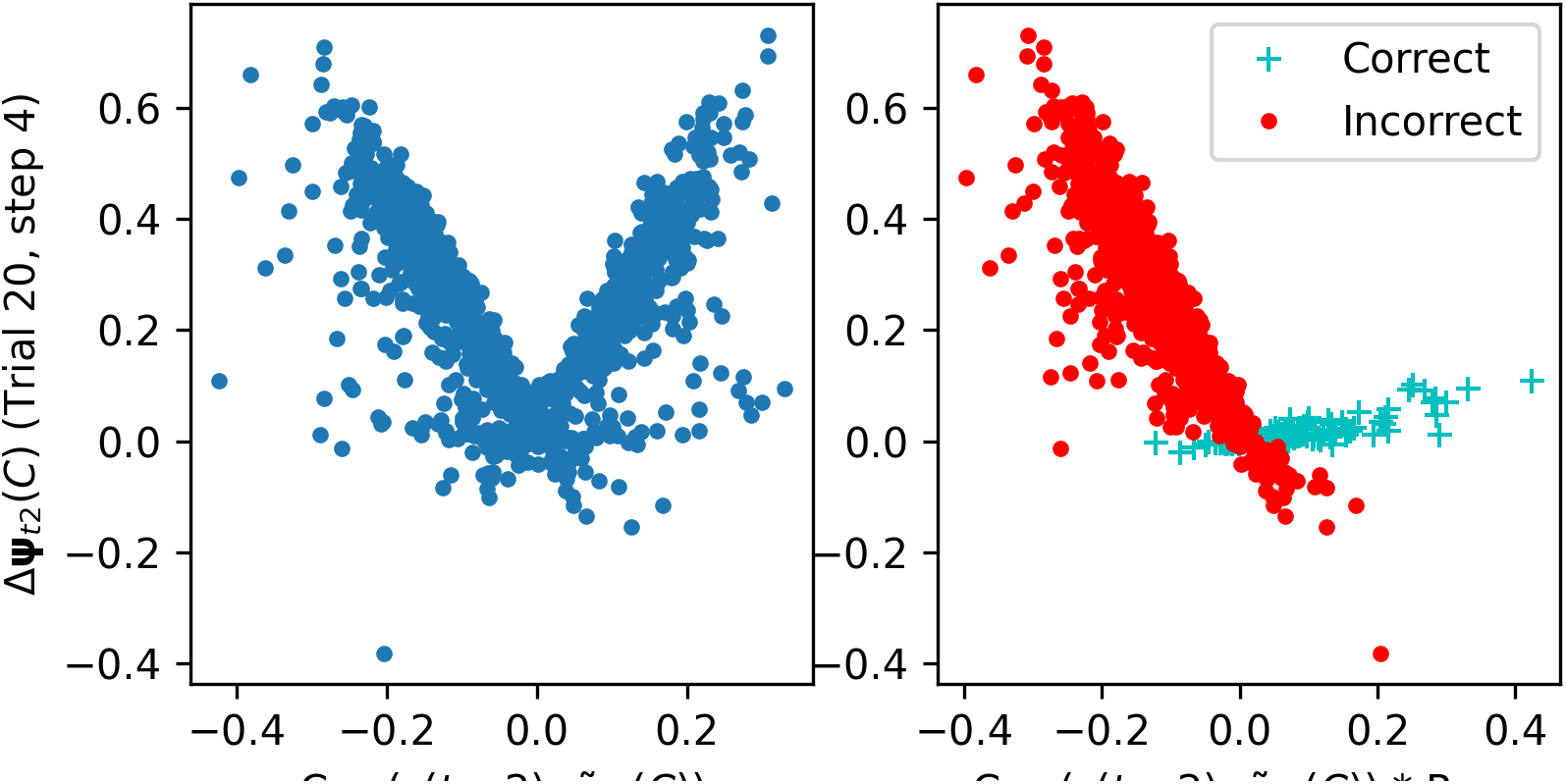
Left: Signed amplitude of reinstated vector 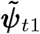 at time step 2 predicts representation change for *C* at time step 4, in trial 20 (with pair *DE* or *ED* shown as stimulus for this trial). Right: Same data but multiplying *x* by the sign of the response given for this trial, with separate coloring for correct and incorrect responses. Data contains 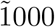 runs with *DE* not shown before trial 20 (leading to incorrect responses) and 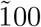 trials in which it was (leading to mostly correct responses).

## Footnotes

1 https://github.com/ThomasMiconi/TransitiveInference

2 If we do not reset neural activations between trials, the network consistently develops strategies that process information from successive trials in parallel; that is, activations in trial *k* represent information not just from trial *k* but also from trial *k −*1. While potentially interesting in itself, this phenomenon complicates the investigation of mechanisms underlying within-episode learning. We therefore chose to reset activations at the start of each trial.

3 For example, suppose we show items *D* and *E* at a given trial. In future trials, if pair *EF* is shown, we want *D* to be reinstated with the same sign as *E*, since both *D* and *E* are “left” of *F*. Similarly, if pair *CD* is shown, we wand *E* to be reinstated with the same sign as *D*, since both *D* and *E* are “right” of *C*.

